# The molecular logic of synaptic wiring at the single cell level

**DOI:** 10.1101/2020.11.30.402057

**Authors:** Jessica Velten, Rashi Agarwal, Patrick van Nierop, Katrin Domsch, Lena Bognar, Malte Paulsen, Lars Velten, Ingrid Lohmann

## Abstract

The correct wiring of neuronal circuits is one of the most complex processes in development, since axons form highly specific connections out of a vast number of possibilities. Circuit structure is genetically determined in vertebrates and invertebrates, but the mechanism guiding each axon to precisely innervate a unique pre-specified target cell is poorly understood. Here, we used single cell genomics, imaging and genetics to show that single-cell specific connections between motoneurons and target muscles are specified through a combinatorial code of immunoglobulin domain proteins. These programs are orchestrated by a homeodomain transcription factor code that specifies cellular identities down to the level of biologically unique cells. Using spatial mapping, we show that this code follows spatial patterns already observed at early embryonic stages, while acting at a much later stage to specify motor circuits. Taken together, our data suggest that a relatively simple homeo-immunoglobulin-code determines neuronal circuit structure.

## INTRODUCTION

Neuronal circuits in the nervous system of mammals as well as in *Drosophila* are stereotypically wired for the precise execution of functional tasks critical for organismal survival. The formation of such circuits is a step-wise process, which starts with the specification of neuronal cell types and their accurate arrangements in space, followed by the correct wiring of individual cells and their final integration into a functional network. To ensure such precision, the structure and connectivity of neural circuits is genetically specified. However, how these complex interconnected processes are encoded in the genome and executed by the cellular protein machinery remains an unresolved question.

According to the ‘labelled pathway hypothesis’ (Sperry, 1963), neurons stochastically and transiently form contacts with many possible targets after their specification, while the expression of specific cell surface proteins (CSPs) is thought to stabilize the correct connections, a process called synaptic specificity. Many lines of evidence support this hypothesis. Recent work has shown that combinations of IgSF cell surface proteins (Dprs) are differentially expressed in distinct neuronal clusters and bind to specific Dpr binding proteins (DIPs) expressed in synaptic partners (Carrillo et al., 2015; Nakamura et al., 2002; Özkan et al., 2013). In the visual system combinations of CSPs are expressed differentially in different layers (Tan et al., 2015), while in olfactory neurons, a combinatorial code of TFs and CSPs maps neurons with the same olfactory receptor to the same glomerulus (Couto et al., 2005; Li et al., 2017, 2020). All these studies explain how groups of similar neuronal cells are molecularly defined and provide a hypothesis on how stereotypic connections to another neuronal cell type are formed. By contrast, how the specificity of circuits is specified at the level of single cells has not been addressed.

In the *Drosophila* neuromuscular system, every single motoneuron within the developing embryo forms unique and stereotypic connections with target muscles, but the molecular mechanism underlying this specificity is unclear (Allan & Thor, 2015). Motoneurons are progressively specified from anterior to posterior by segment specific TFs (Angelini & Kaufman, 2005; Bossing et al., 1996; Schmid et al., 1999; Schmidt et al., 1997), and further along the dorsal to ventral axis (Broihier et al., 2004; Broihier & Skeath, 2002; Matthias Landgraf & Thor, 2006) before extending their nerve projections to predefined locations specified in all three spatial dimensions (Broihier et al., 2004; Broihier & Skeath, 2002; Hessinger et al., 2017; Matthias Landgraf et al., 1997; Thor et al., 1999). These observations have suggested region-specific mechanisms in the determination of connectivity patterns. However, such a regional model alone is unlikely to explain the precise connectivity patterns of single cells (Matthias Landgraf et al., 2003; Nassif et al., 1998). Elegant studies in the vertebrate central nervous system and classical transplantation experiments demonstrated in addition that positional identity and connectivity patterns of single neurons are stably maintained even after experimental relocation of cells (Demireva et al., 2011). Thus, there is apparently a molecular mechanism that stably imprints cellular identity and instructs the formation of connectivity.

Homeodomain transcription factors (TFs) have long been known to play important roles in the specification and differentiation of neurons (Allen et al., 2020; Deneris & Hobert, 2014; Domsch et al., 2019; Reilly et al., 2020; Sugino et al., 2019; Urbach et al., 2006, 2016; Zeisel et al., 2018). Importantly, it has been shown just recently that each neuron class in the nematode *C. elegans* expresses a unique combination of homeodomain TFs thereby unambiguously specifying neuronal identities (Reilly et al., 2020). However, it is so far unclear whether this concept extends to other organisms; if such a cell specific homeo-code also instructs later events in circuit formation; and, finally, which molecules downstream of such a TF code realize synaptic target choice and specificity at the single cell level.

Using single cell RNA sequencing (scRNA-Seq) with high numbers of biological replicates we demonstrate the existence of a homeo-code defining the cellular identities and positions of differentiated *Drosophila* motoneurons along the major body axes. We exploit this code to show that a highly specific map of cell surface proteins (CSP), in particular cell surface Immunoglobulins (Igs), determines specificity during the synaptic wiring phase. Knockdown and ectopic expression of homeodomain TFs induces synaptic wiring defects specific to single cells that are phenocopied by genetic modulations of their Ig targets. Additionally, we demonstrate that common homeo-code signatures label matching synaptic partners of functional neuronal circuits. In sum, our data reveal how development of individual neuronal circuits is genetically specified by a relatively simple and linked ‘homeo-immunoglobulin program’, determining complex neuronal wiring with single cell precision.

## RESULTS

### A reference map of motoneurons during the synaptic wiring phase in *Drosophila* embryos

To generate a spatially resolved molecular map of motoneurons during the synaptic wiring phase at the end of embryogenesis, we performed scRNA-Seq of stage 17 embryonic cells marked by the motoneuron-specific *OK371-*GAL4 driver (Mahr & Aberle, 2006) controlling the expression of the UAS-*RFP* transgene. We used this experimental design for the following reasons: first, the glutamatergic driver line (*OK371-*GAL4) is based on a regulatory element controlling the glutamatergic neurotransmitter receptor (VGlut), a gene which is expressed in motoneurons when synaptic connections are established; and second, it has been shown before that neurons diversify most during the synaptic wiring phase and become indistinguishable upon completion of neuronal connectivity (Li et al., 2017). Thus, we profiled motoneuron transcriptomes during a very narrow one-hour time-frame at the end of embryogenesis when the first connections between motoneurons and muscles are formed (Fig. 1A).

**Figure 1:**
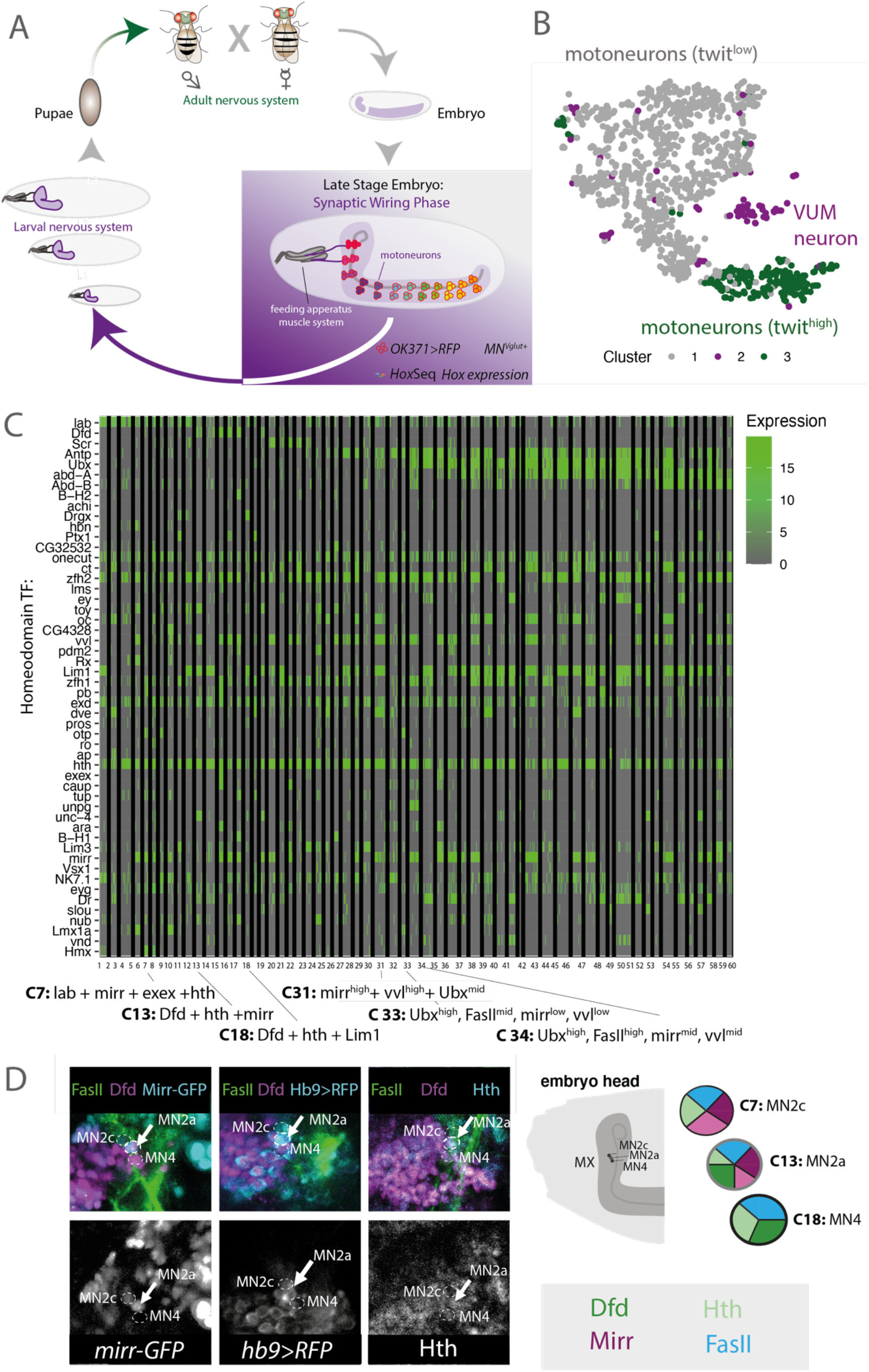
scRNA-Seq identifies anterior-to-posterior regional variability as a major determinant of motoneuronal heterogeneity. **(A)** Schematic drawing depicts the development of distinct neuronal systems during *Drosophila* lifetime. The first round of synaptic wiring in the late *Drosophila* embryo is highlighted and depicts the lateral view of an early stage 17 *Drosophila* embryo. Experimental design: Motoneurons expressing UAS-*RFP* under control of *OK371-*GAL4 are color-coded along the ventral nerve cord according to different patterns of *Hox* gene expression. scRNA-Seq protocol of motoneurons was performed using a targeted protocol to enrich for *Hox* gene representation (HoxSeq). **(B)** T-distributed stochastic neighbor embedding (t-SNE) plot of n=999 single-cell transcriptomes. Colors correspond to clusters identified using hierarchical clustering that were annotated using marker gene expression as shown in Figure S1C-S1E. **(C)** Heatmap depicting the expression of homeodomain encoding genes (rows) across n= 758 single *twit^low^* motoneurons (columns). Rows and columns are arranged by hierarchical clustering. Normalized expression levels are color-coded. Arrows indicate selected clusters for follow up studies shown in Figure 1D and Supplementary Fig. 3B. **(D)** Visualization of homeodomain TFs by immunohistochemistry in the head of a stage 17 *Drosophila* embryo, lateral view. Pie charts represent single cells with corresponding homeodomain TF expression (color code). The combinatorial expression of the four homeodomain TFs Dfd, Mirr (*mirr-GFP*), Exex (*Hb9>RFP*) and Hth is exclusive to the MN2a motoneuron (FasII^+^). The homeodomain TF Dfd is absent in motoneuron A (MN2c), while the homeodomain TFs Mirr and Exex are absent in MN4.

Development of synaptic connections in the *Drosophila* nervous system is a step-wise process, beginning with a low number of stable synaptic connections in late embryos and a significant increase in the number of connections throughout larval development (Couton et al., 2015; Mahr & Aberle, 2006). In this study, we focused on the first phase of synaptic wiring at the end of embryogenesis, therefore a significant lower number of *OK371*-positive motoneurons interacting with a target muscle is expected in comparison to the final set of 35 motoneurons in each hemisegment of the larval nervous system. Using microscopy, we identified approximately 140 cells in total to be positive for *OK371* activity during stage 16 and early stage 17 (Supplementary Fig. 1A). It is this low number of glutamatergic motoneurons in early wiring, which allowed for a high number of biological replicates of biologically unique cells in scRNA-Seq. Single *RFP*-expressing motoneurons were sorted from a pool of precisely staged embryos by Fluorescence-Activated Cell Sorting (FACS) (Fig. 1A). In total, 1536 motoneurons were sequenced by SMART-Seq2 (Picelli et al., 2014). After filtering, 999 single-cell transcriptomes were retained, thus each biologically unique motoneuronal cell was covered on average 7-fold in our dataset (Supplementary Fig. 1B). Abundant expression of motoneuronal marker genes like *Vesicular glutamate transporter (VGlut)* and *embryonic lethal abnormal vision (elav)* indicated successful sorting of the targeted cell population (Supplementary Fig. 1D). A median of 1202 genes were observed per cell (Supplementary Fig. 1B), and a negligible fraction of 0.1% of reads mapped to the mitochondrial genome, supporting the high technical quality of the data.

*Hox* genes are known to be expressed in a consecutive order along the anterior-posterior (AP) of *Drosophila.* We used this property to precisely locate single cell transcriptomes along the AP axis as further described below. To this end, we implemented a custom modification of the SMART-Seq2 protocol by adding primers targeting each *Hox* gene to the reverse transcription (RT) and preamplification step that permits an increased representation of *Hox* gene expression as spatial markers (Fig. 1A, see Materials and Methods). Despite the low expression of *Hox* genes in late embryonic stages, we identified 75% of the motoneurons to express at least one *Hox* gene (Supplementary Fig. 1E).

To explore the molecular diversity of the motoneurons, we performed two independent unsupervised analyses, t-distributed neighbor embedding (tSNE) and hierarchical clustering (Fig. 1B, Supplementary Fig. 1C). Both methods identified a cluster corresponding to modulator neurons (VUMs, 8% of the cells) (Fig. 1B, Supplementary Figs 1C, 1D, 1F) as well as two large yet distinct clusters of motoneurons that differ in the expression of the marker genes *rhea* and *target of wit* (*twit*) (Supplementary Fig. 1F). VUM motoneurons belong to a very distinct motoneuron subtype expressing a combination of subtype specific marker genes, *Vesicular monoamine transporter (Vmat), Tyramine β* hydroxylase (*Tbh), diacyl glycerol kinase (dgk)* and the motoneuronal marker *Zn finger homeodomain 1* (*zfh1)* (Stagg et al., 2011a), which we all identified in our modulator neuron cluster. These type II glutamatergic/octopaminergic motoneurons exhibit modulator roles in taste responses (Matthias Landgraf et al., 1997; Siegler & Jia, 1999; Sink & Whitington, 1991; Stagg et al., 2011b), while the *twit^l^*^ow^ and *twit*^high^ cluster can be assigned to the abundant glutamatergic type I motoneuron class (Hoang & Chiba, 2001; M. D. Kim et al., 2009). *In situ* hybridizations of late stage embryos localized *twit* transcripts in median and lateral clusters of posteriorly located motoneurons (N. C. Kim & Marqués, 2012), suggesting that those two groups of motoneurons represent indeed two distinct motoneuronal subtypes with different locations.

Although the motoneuronal population in the largest cluster (*twit*^high^) appears rather homogenous, our data set confirms mutual exclusive expression of known marker genes expressed in subsets of dorsally and ventrally projecting motoneurons (Garces & Thor, 2006a; Kania & Jessell, 2003; M Landgraf et al., 1999). In particular, the dorsal marker gene *Drop (Dr/msh)* and the ventral marker genes *Hb9/exex* are expressed in distinct populations (Couton et al., 2015). Likewise, the dorsal marker gene *grainy head (grn)* or *LIM homeobox 1 (Lim1)* (Kania & Jessell, 2003) and the ventral marker gene *islet (isl/tup)* (Garces & Thor, 2006b) are expressed in different subsets (Supplementary Fig. 1G).

Taken together, these analyses showed that the dataset generated in this study was of high quality, consistent with published data and provided an approximately 7-fold cellular coverage of each biologically unique MN at the synaptic wiring stage.

### A homeodomain TF code specifies single *Drosophila* motoneurons during synaptic wiring

To investigate processes required for motoneuron diversity during the synaptic wiring phase, we focused on the largest cluster of motoneurons expressing low levels of *twit* (*twit^low^*). Our reasoning was that this cluster contains the majority of glutamatergic type I motoneurons that are equally distributed along the VNC, rather than specialized subtypes of motoneurons such as the VUMs or *twit*^high^ clusters that are unevenly distributed along this embryonic body axis. This is particularly important, because in this analysis we focused on processes required for synaptic wiring rather than on molecular programs distinguishing cell subtype identities. As a first step, we used unsupervised identification of highly variable genes and principal component analysis (PCA) of the *twit*^low^ cells and identified variability in several biological processes, most notably homeodomain TF expression as well as the expression of genes encoding CSPs involved in synaptic matching and axon guidance (Supplementary Fig 2A-E). In particular, the expression of known mediators of synaptic specificity, for instance Dpr protein encoding genes (Carrillo et al., 2015), was highly variable within the *twit^low^* cluster, and co-varied with homeodomain TF gene expression (Supplementary Figs 2A-E).

Using a detailed analysis of homeodomain TF expression, we observed that small groups of cells were defined by a unique expression pattern, an expression ‘code’ (Fig. 1C). These patterns were independent of technical covariates (Supplementary Figure 2F). In our experimental design, we focused in particular on the subset of type I notoneurons (*twit^l^*^ow^ cluster) whilst each biologically unique cell was covered approximately 7 times, raising the possibility that unique homeodomain TF expression patterns correspond to symmetrical pairs of motoneurons with biologically unique identities (120 unique single cells, two of which share the same molecular identity due to the bilateral symmetry, for a total of 60 unique molecular states). These findings corresponds to a recently described homeodomain ‘code’ described in single neuron classes of *C. elegans* (Reilly et al., 2020). Since a statistic specification of cluster number from scRNA-Seq data remains an unresolved issue in the field (Luecken & Theis, 2019; Zhu et al., 2019), we used immunofluorescence stainings of homeodomain TFs in motoneurons to validate the single-cell specificity for six out of the 60 clusters from the observed code. This included a motoneuron, which according to our scRNA-Seq data was suggested to co-express Deformed (Dfd), Homothorax (Hth) and Mirror (Mirr) (Fig. 1C). This neuron should be located in the Dfd+ region (i.e. the maxillary segment of the AP axis) and in the ventral region along the DV axis (Mirr). In fact, the co-expression of these factors was confirmed in a motoneuron, which projects via the maxillary nerve to the mouth hook elevator muscle (MHE) and emerges ventrally to the antennal nerve of the maxillary segment (Fig. 1D, Supplementary Fig. 3A). Based on its similar position in *Calliphora vicina*, we termed this axon projection MN2a (Schoofs et al., 2010). Another motoneuron coexpresses Dfd and Hth, but is lacking the expression of the ventral markers Mirr and projects to the dorsal bundle of the mouth hook depressor (MHD) in larvae (termed MN4 based on (Schoofs et al., 2010). We also validated the MN2c motoneuron, which co-expresses the same set of genes as MN2a but lacks the segment defining Dfd expression (Fig. 1D). Thus, it projects anterior to the labial retractor (LR) muscle in larvae. As further examples of homeodomain co-expression patterns, we confirmed the existence of a motoneuron originating from the A2 segment with intermediated (mid) levels of Ultrabithorax (Ubx) and FasII and high levels of Mirr and Vvl, a single motoneuron with high levels of Ubx, intermediate levels of FasII and low levels of Mirr and Vvl (Supplementary Fig. 3B), as well as a single motoneuron with high Ubx and FasII levels and intermediate Mirr and Vvl levels. Thereby, we confirmed the existence of six individual motoneurons by immunofluorescence stainings corresponding to clusters C7, C13, C18, C31, C33 and C34 in our scRNA-Seq data set. In sum, these analyses revealed a homeodomain TF ‘code’ as unique identifier of cellular identity during the synaptic wiring phase.

### Regional specific expression of the homeo-code

The identification of a homeo-code raised the possibility that cells possess a molecular memory of their spatial position that later coordinates synaptic wiring. To investigate this hypothesis, we mapped single motoneurons in space. For mapping neurons along the AP axis, we created a high-resolution map of Hox protein expression using immunofluorescence (Fig. 2A, left panel, Supplementary Figs 4A, 4B). Anterior Hox proteins were expressed in clearly defined stripes, whereas posterior Hox protein expression patterns partly overlapped. The same co-expression patterns were observed in our scRNA-Seq data (Fig. 2A, right panel), allowing us to probabilistically map each cell from the scRNA-Seq data set to a position along the AP axis (Fig. 2A, Supplementary Fig. 4C, Materials and Methods). We validated this mapping strategy by immunofluorescence. To this end, we used the inferred AP position to estimate the expression pattern of every gene along the AP axis (Fig. 2B). We thereby identified candidate genes with differential expression along this axis. Importantly, these candidates were not used for constructing the model. For two such candidates, *Frq1* and *hth*, we compared the predicted expression pattern to immunofluorescence data and observed a high agreement (Supplementary Fig. 4D). Inferred AP position was highly correlated with principal component 3 and 4, indicating that AP position profoundly affects the entire transcriptome of the cell (Supplementary Fig. 4E). More importantly, the above described homeo-code is aligned with the AP position, but on the fine scale the homeo-code shows more variability independent of AP position (Supplementary Fig. 4F). In order to identify these additional processes, we used ZINB-WaVE (Risso et al. 2018) to separate scRNA-Seq data into variability linked to the known covariates (AP position and technical variability), and processes statistically independent thereof (Fig. 2C). Thereby, we identified on the first component of AP-independent variability one group of genes known as marker for dorsal-ventral (DV) position (Bhat, 1999; Skeath, 1999; Urbach et al., 2006). These genes were ordered according to their localization in the embryo from dorsal to ventral (Fig. 2D). Again, homeodomain TFs and Ig surface proteins were among the most variable genes on this AP-independent axis of variability (Fig. 2E). Immunofluorescence experiments of two ventral marker genes, *mirror (mirr)* and *ventral veins lacking (vvl),* confirmed the predicted DV position (Fig. 2F, Supplementary Fig. 4G).

**Figure 2:**
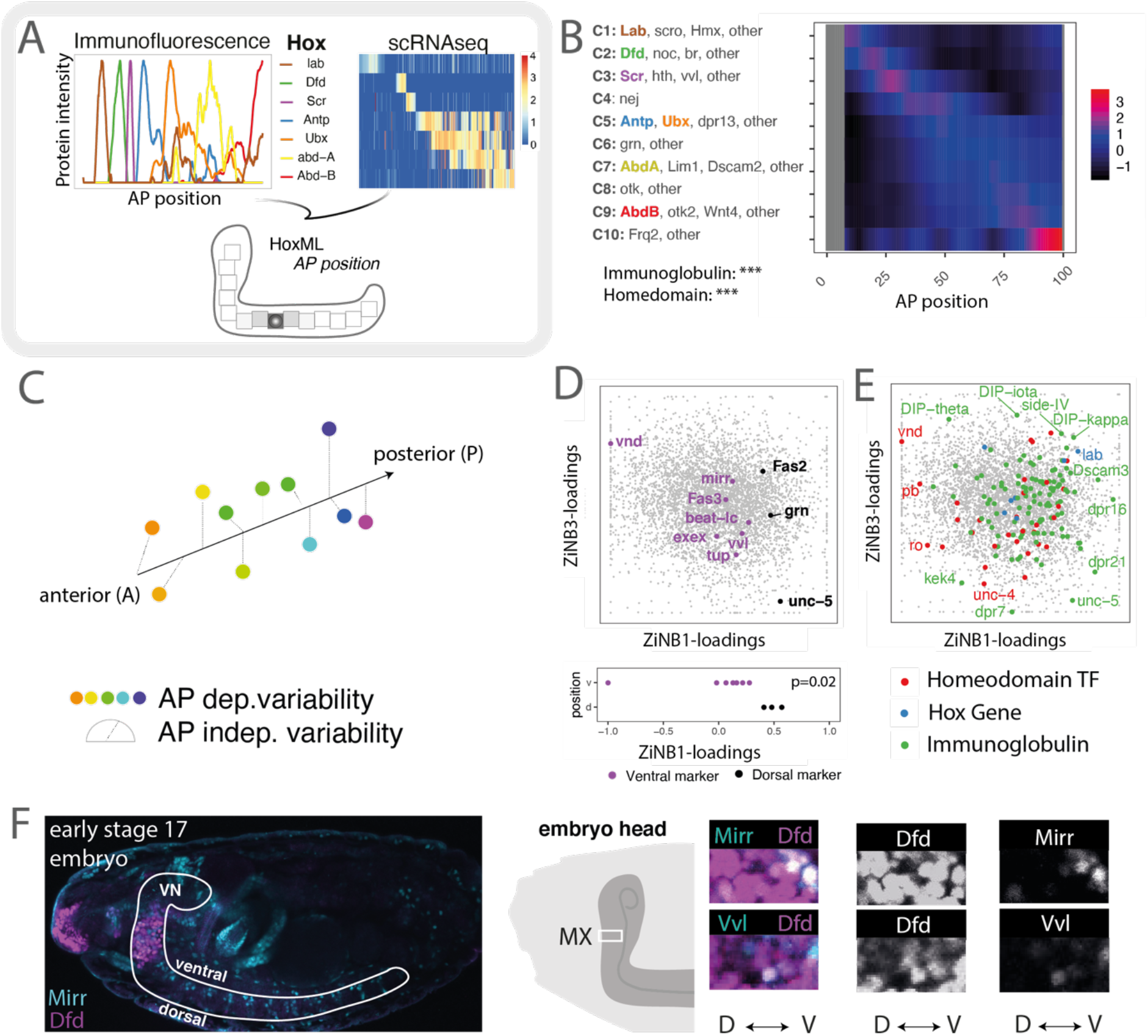
Unsupervised analysis of scRNA-Seq data identifies single-cell specificity of homeodomain gene expression patterns. **(A)** Outline of strategy to map single motoneurons to position along the AP axis. *Left panel:* Intensities of Hox protein expression along the ventral nerve cord measured by immunofluorescence, see also Figure S3A, S3B. *Right panel:* Co-expression pattern of *Hox* gene transcripts measured by scRNA-Seq, see also Figure S3C. Expression level is color-coded; columns correspond to n=758 single *twit^low^* motoneurons. *Bottom panel:* AP position is inferred form scRNA-Seq data by probabilistically mapping *Hox* gene expression pattern in each individual cell to the immunofluorescence reference data. See Materials and Methods. **(B)** Genes with significant variation along the AP axis were identified and clustered into 10 groups of distinct expression pattern (Materials and Methods). Heatmap shows average gene expression per cluster (rows) across single cells (columns). See Supplementary Table 1 for a complete list of genes in each cluster. Asterisks indicate p-value of a hypergeometric test for enrichment of protein domains, ***: p < 0.001. **(C)** ZiNB-WaVE (Risso et al., 2019) was used to statistically separate gene expression variability into parts linked to AP position and parts independent thereof. **(D)** Scatter plot of ZINB-WaVE loadings separates known dorsal and ventral marker genes on ZiNB-WAVE component 1. **(E)** Genes encoding homeodomain TFs and genes encoding Ig domain molecules (see color code) show high loadings on ZINB-WaVE component 1 and 3, demonstrating high variability independent of AP position. **(F)** Representative confocal picture of an early stage 17 *Drosophila* embryo, lateral view. Dorsal and ventral site of elongated ventral nerve cord (VN) are highlighted. Visualization of two ventral homeodomain TF markers (Mirr, Vvl) in the maxillary segment (MX).

Together, these analyses suggested that the position along the animal body axis determines the homeo-code and thereby defines cellular identity.

### Homeodomain TFs modulate synaptic target specificity in a position dependent manner

Are homeodomain TFs functionally linked to processes mediating synaptic specificity? It is described that this class of TFs is involved in maintaining the neuronal lineage (Deneris & Hobert, 2014; Domsch et al., 2019; Friedrich et al., 2016; Reilly et al., 2020). However, it is so far unclear whether sets of homeodomain TFs instruct specific programs for precise matching of motoneurons and their targets. To address this question, we interfered with *Dfd*, *mirr* and *hth* by RNAi using the pan-neural driver *elav-*GAL4 (Luo et al., 1994) and examined synaptic defects in early stage 17 *Drosophila* embryos. Despite the activity of the driver in neuroblast stages, RNAi mediated knockdown is realized earliest in stage 13-14 of motoneuronal axonogenesis (Supplementary Figs 5A, 5B, 5C). Thus, the driver is ideally suited for temporal interference of proteins during the synaptic wiring phase. To complement this approach, we used in addition a more specific driver, *OK6-*GAL4, which is active in glutamatergic motoneurons only (Mahr & Aberle, 2006).

Defects induced by RNAi mediated gene silencing were classified into two distinct categories, wiring defects (i.e. mistargeting of neurons to the wrong muscle), and terminal defects (i.e. abnormal synaptic morphologies at axon terminal sites). We focused our analyses on the MN2a (Fig. 2F), since this motoneuron can be easily identified by the co-expression of Dfd and FasII. MN2a innervates the mouth hook elevator muscle (MHE), an interior muscle of the *Drosophila* feeding apparatus that also controls hatching movements (Fig. 3A) (Friedrich et al. 2016). Thus, an assay of hatching rates is a relatively simple readout to examine the impact of wiring defects on behaviour. Embryos depleted for *Dfd*, *mirr* and *hth* were all characterized by significant morphological changes of the MN2a motoneuron (Fig. 3B). 70% of *hth* depleted embryos displayed severe wiring defects, including mistargeting of the MN2a to ventral and anterior muscles of the outer skeleton of *Drosophila* embryos (Figs 3B, 3C), while 29% of the *Dfd* depleted embryos showed mistargeting of MN2a to the Labial Retractor muscle (LR) located directly anterior to the MHE in the Lab^+^, Dfd^-^ segment (Figs 3B, 3C), leading to a decreased hatching rate (Fig. 3D). Survivors of the hatching rate assay showed proper targeting of MN2a to the MHE, but abnormal synaptic morphologies, such as an increased number of synapses (Figs 3B, 3E). In the case of *mirr* knockdown, wiring effects were less pronounced and frequent. Together, these results suggested that loss of homeodomain TF expression leads to synaptic targeting defects.

**Figure 3:**
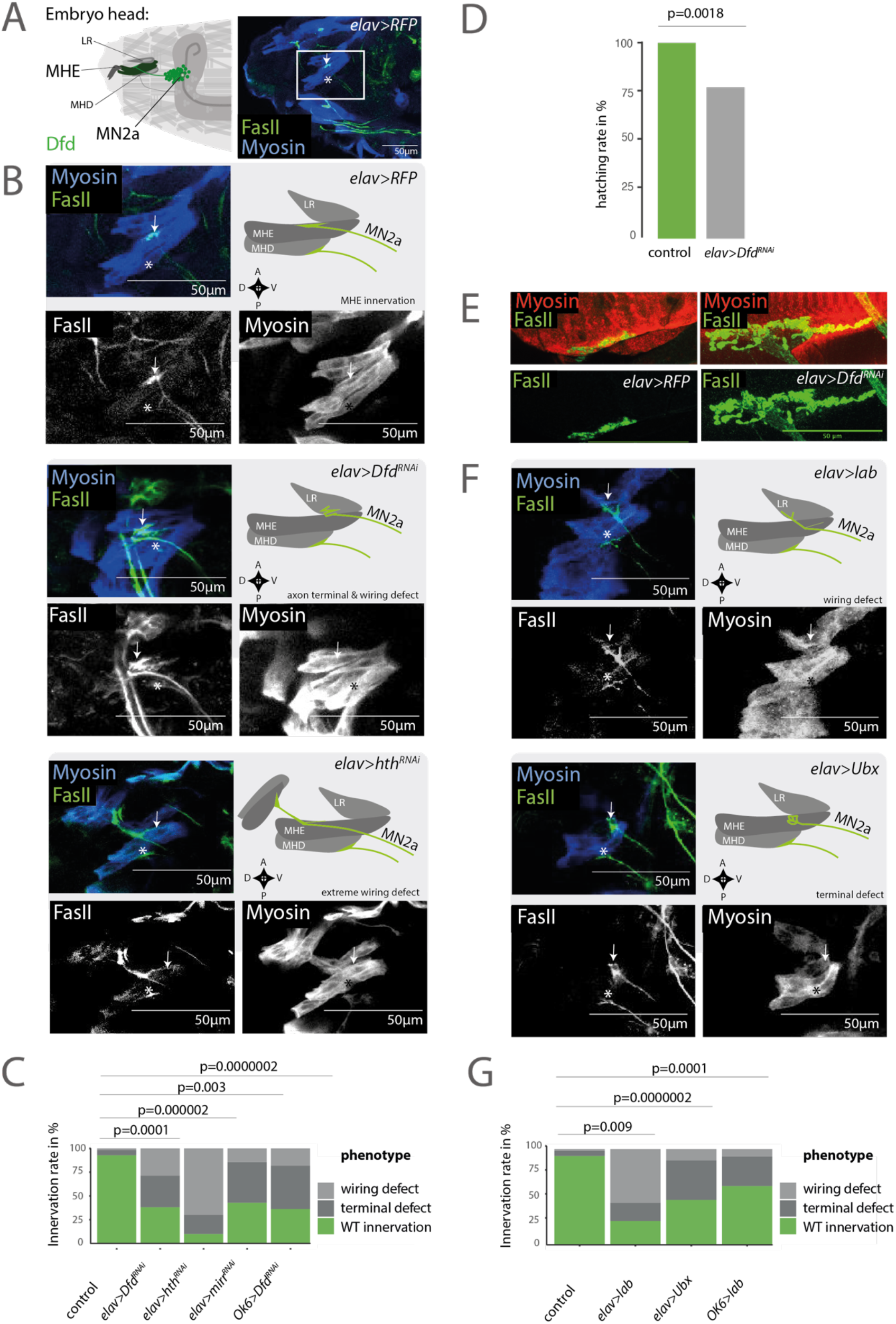
Homeodomain TFs modulate target specificity of motoneurons. **(A)** *Left panel:* Schematic drawing depicts lateral view of an early stage 17 *Drosophila* embryonic head. The MN2a motoneuron expressing Dfd (light green) innervates the MHE (dark green). The MHE controls elevation movements of mouth hooks (dark grey) that are required for hatching from the eggshell (Friedrich et al., 2016). *Right panel:* Representative confocal picture of an early stage 17 *Drosophila* embryonic head depicts the innervation site (white arrow) on the MHE muscle (blue) by MN2a (green). The innervation site on the MHD muscle (blue) is highlighted by an asterisk as reference for orientation in the embryo. **(B)** Visualization of three independent genetic experiments using the pan-neural *elav*-GAL4 driver to control the expression of UAS-*RNAi* constructs of homeodomain TFs i. e. UAS-*Dfd^RNAi^*, UAS-*hth^RNAi^* and UAS-*RFP* as control. *Left panels:* Zoom on MHE of an early stage 17 *Drosophila* embryo. Note the upper panel shows a zoom of the image depicted in Figure 3A as control. *Right panels:* Schematic drawing depicts the innervation phenotype on MHE by the MN2a motoneuron. Phenotypes are classified for abnormal synaptic morphologies at axon terminal sites: ‘terminal defects’ and incorrect innervation of target muscle ‘wiring defects’. In wild-type conditions, the MN2a motoneuron projects to the MHE, while in *elav>Dfd^RNAi^* animals MN2a motoneurons display axon ‘terminal defects’ and ‘wiring defects’, i.e. incorrect targeting to the LR muscle and in *elav>hth^RNAi^* animals incorrect targeting of an outer skeleton muscle, termed ‘extreme wiring defects’. **(C)** Phenotypic penetrance for MN2a projections ‘innervation rate’ was calculated by the rate of correct axon projections of MN2a to MHE (mock; n=56) compared to abnormal synaptic morphologies at axon terminals ‘terminal defects’ and wiring defects displayed by animals expressing RNAi constructs for homeodomain TFs i. e. *OK6>Dfd^RNAi^* (n=11), *elav>Dfd^RNAi^* (n=21), *elav>mirr^RNAi^* (n=7) and *elav>hth^RNAi^* (n=10). Note, each genetic experiment was performed in parallel to an adequate control experiment using the same driver line crossed to a line that controls expression of either UAS-*RFP* or UAS-*GFP^RNAi^*. For quantifications these control experiments were summarized and termed ‘mock’ experiments. p-values between two genetic conditions were calculated by a two-sided hypergeometric test. **(D)** Correct MHE innervation is required for hatching of *Drosophila* embryos from the eggshell (Friedrich et al., 2016). Hatching rate was calculated based on the number of L1 larvae observed after 24h in genetic crosses, depleted of Dfd (UAS-*Dfd^RNAi^*) in neurons (*elav-*GAL4; n=217) compared to crosses with control animals (mock = *elav>RFP*; n=156). **(E)** Representative confocal picture depicts the innervation site on the MHE muscle in L3 larvae. Animals depleted of Dfd in neurons (*elav>Dfd^RNAi^*) show enlarged innervation sites and increased amounts of boutons and branching phenotypes (right panel) compared to control animals (*elav>RFP*), (left panel). **(F)** *Right panel:* Representative confocal images of two independent genetic experiments using the pan-neural *elav*-GAL4 driver to control the expression of the homeodomain TFs Ubx and Lab i. e. UAS*-Ubx,* UAS*-lab*. The control experiment (*elav>RFP*) is depicted in Figure 3B. *Left panel:* Zoom on MHE of an early stage 17 *Drosophila* embryo. *Right panel:* Schematic drawing depicts the innervation phenotype on MHE by the MN2a motoneuron. See Figure 3B for classification of phenotypes. In wild-type conditions the MN2a motoneuron projects to the MHE, while in *elav>lab* animals MN2a motoneurons display ‘wiring defects’, i.e. incorrect targeting to the LR muscle and in *elav>Ubx* animals show aberrant morphologies at axon terminals, termed ‘terminal defects’. **(G)** Innervation rates of *OK6>lab* (n=13), *elav>lab* (n=16) and *elav>Ubx* (n=17) animals are calculated as described in Figure 3C.

The hypothesis of a homeodomain ‘code’ implies that not only the loss of factors but also their ectopic expression should cause wiring defects. We therefore ectopically expressed the *Hox* genes *labial* (*lab)* as well as *Ubx* in all neurons using the *elav-*GAL4 driver. Lab is expressed directly anterior to Dfd, while Ubx is active in abdominal segments. Pan-neural and motoneuron-specific misexpression of *lab* led to mistargeting of MN2a to the LR muscle (Supplementary Fig. 5D), reminiscent of the mistargeting phenotype observed in *Dfd* knockdown conditions (Figs 3F, 3G). By contrast, *Ubx* mis-expression typically elicited terminal defects but not a wiring phenotype. In sum, stage specific genetic interference with homeodomain TFs resulted in changes in target specificity of motoneurons, supporting the idea that these factors modulate synaptic specificity in late embryonic stages when synaptic connections are established.

### Homeodomain effects on target specificity are mediated by combinatorial immunoglobulin expression

Our single-cell analysis and functional follow-ups revealed a critical role for homeodomain TFs in controlling synaptic specificity in the neuromuscular system, motivating an investigation of potential downstream effectors. Unsupervised analysis of gene classes associated with the homeo-code revealed that Ig encoding genes most strongly correlated with homeodomain clusters (Figs 4A, 4B, 4C see also Supplementary Figs 2D, 2E). More detailed analysis allowed us to predict specific Ig codes (Fig. 4C, see also Supplementary Fig. 5F-H) for every homeodomain cluster. Thus, our data set provides a comprehensive overview of the distribution of Ig encoding genes across different cellular identities. For example, segment specific expression of Ig encoding genes is observed for *off-track 2 (otk2)*, a gene enriched in posterior segments. Some other Ig encoding genes, such as those encoding Down syndrome cell adhesion molecules (Dscams) or Kekkon (Kek) neurotrophin receptors (Ulian-Benitez et al., 2017), are expressed in most homeodomain clusters, but are occasionally switched off. This phenomenon we term here “OFF code”. In other cases, such as genes of the DIP family, the *Ig* gene is specifically expressed in one or few homeodomain clusters, which we therefore termed the “ON code”. This code is the most specific Ig code observed in our data set. Together these results indicate that homeodomain TFs directly regulate Ig domain expression. To provide further evidence for the hypothesis that a specific Ig code is controlled by unique combinations of homeodomain TFs, we analysed previously generated whole-embryo Dfd ChiP-Seq data (Sorge et al., 2012) and Ubx ChiP-Seq data, which were retrieved from embryonic neuronal tissues of late stage embryos (Domsch et al., 2019), and found that Ig encoding genes expressed in motoneurons were enriched among the Dfd and Ubx bound targets (Fig. 4D). In addition, homeodomain TF binding sites were overrepresented within regulatory sequences associated with Ig encoding genes (Fig. 4E), strongly supporting a direct regulation of Ig expression by homeodomain TFs (Fig. 4F).

**Figure 4:**
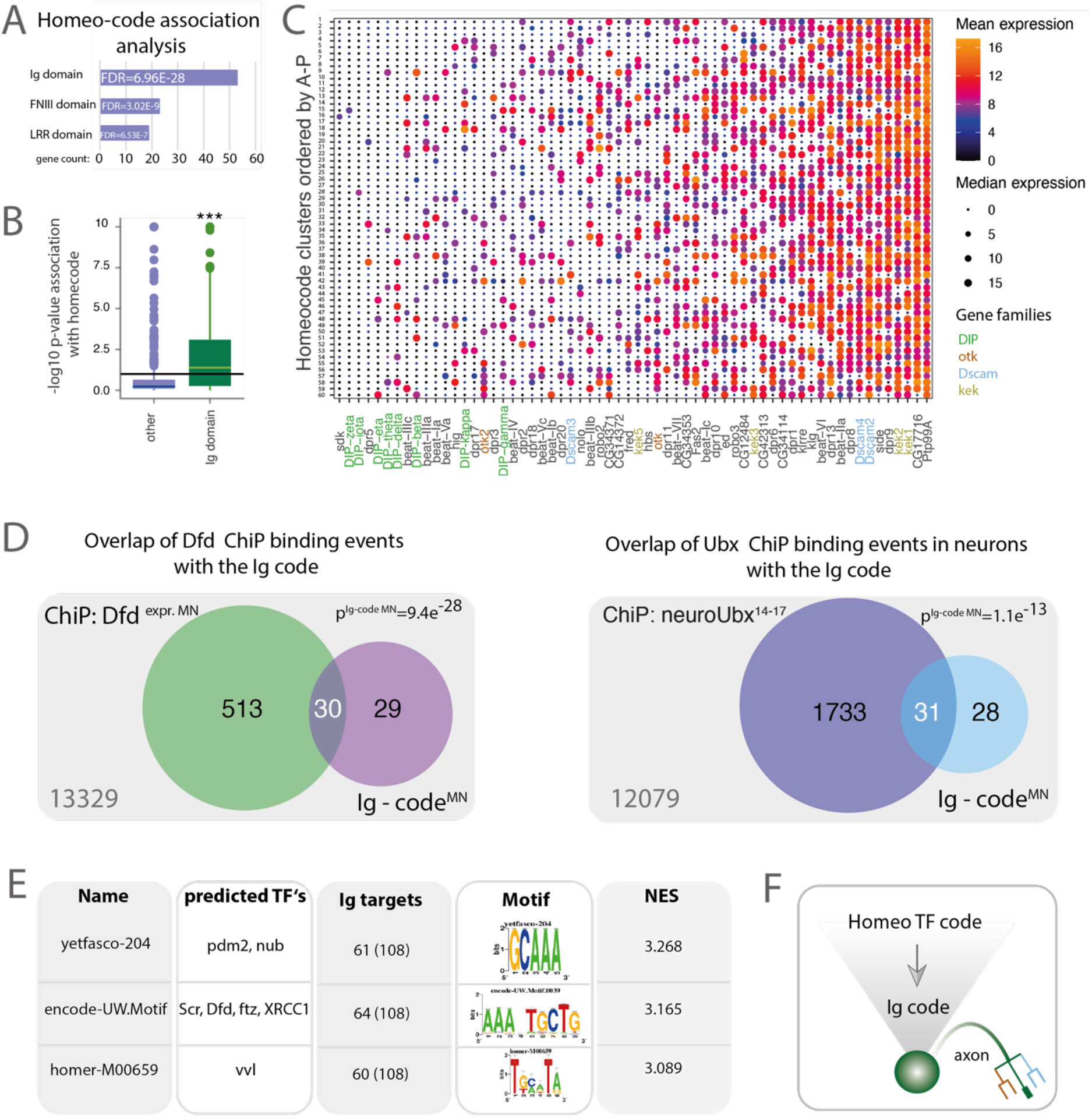
Homeodomain TFs regulate Immunoglobulin (Ig) expression. **(A, B)** Investigation of gene classes associated with the homeo-code. For each gene in the dataset, normalised, scaled expression in single cells was modelled as a function of homeo-code cluster identity, and significance of the association was determined using an F-test. Panel (A) depicts the number of genes with significant homeo-code association falling into distinct gene classes. P values are from a Fisher test. Panel (B) contrasts the P values for homeo-code association between Ig domain genes and other genes using a boxplot. **(C)** Single cells were grouped into 60 clusters according to their expression of genes encoding homeodomain proteins (see Figure 1C) and arranged along the inferred AP position (see Figure 2A). For each cluster, the mean expression of Ig encoding genes was computed. Heatmap depicts Ig encoding genes in columns and clusters in rows; mean expression is color-coded and median expression level is visualized by circle size. Gene families are highlighted in color codes. **(D)** *Left panel:* Venn diagram displays the enrichment of Ig domain genes expressed in motoneurons and bound by Dfd based on the Dfd in a whole embryo ChIP (Sorge et al., 2012). p-value was calculated using a hypergeometric test, n=13920 genes (all protein coding genes expressed in the *Drosophila* genome). *Right panel:* Venn diagram displays the enrichment of Ig domain genes expressed in motoneurons and bound by Ubx in a neuronal tissue specific ChIP (Domsch et al., 2018), n=13920 genes (all protein coding genes expressed in the *Drosophila* genome). **(E)** iRegulon analysis was used to identify TF motifs enriched in the vicinity of Ig encoding genes (see Materials and Methods). 3 of the top 15 highest-ranked motifs of TFs are shown; the predicted targets of these motifs are homeodomain TFs. **(F)** Schematic drawing of the relationship between the homeo-code and the Ig-code in a single neuron.

To investigate the functional role of Ig domain proteins, we focused on the Dfd^+^Hth^+^Mirr^+^ MN2a motoneuron, which highly expressed the Ig-domain proteins DIP-k and DIP-y (Supplementary Fig. 5F). RNAi-mediated knockdown of *DIP-k* in neuronal cells caused both mistargeting of the LR muscle (6%) and terminal defects (41%) with an increased number of synapses formed between MN2a and the target muscle (Figs 5A, 5B, 5C, 5E), similar to the phenotype observed in *Dfd* knockdown embryos (Figs 3B, 3E). In addition, we analysed the hatching behaviour of embryos depleted for *DIP-k* or *DIP-y* and found that their hatching rates were decreased by 93% and 26% respectively when compared to control animals (Fig. 5F). This result indicated that the observed wiring defects are sufficient to impair motor behaviour. Next, we examined the effects of Ig protein mis-expression in neurons on synaptic target selection. A member of the Dpr network, Dpr1, is normally not expressed in MN2a motoneurons according to our single cell data (Supplementary Fig. 5E). Interestingly, misexpression of *Dpr1* in Mn2a resulted in both terminal defects (22%) and wiring defects (22%) such as ectopic targeting of the LR muscle, indicating an altered preference in target choice (Figs 5D, 5E).

**Figure 5:**
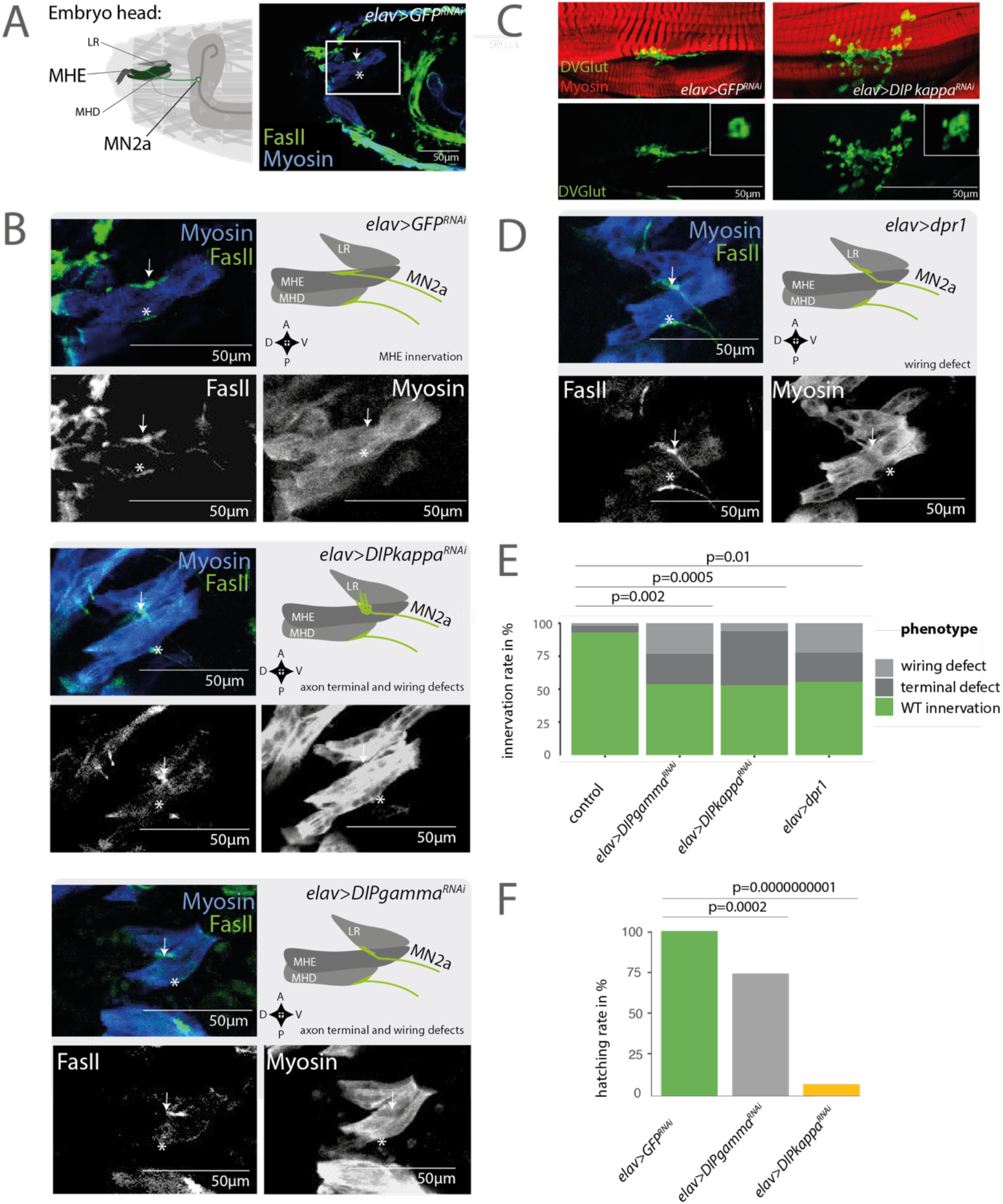
Immunoglobulin proteins are effectors of synaptic specificity. **(A)** *Left panel:* Schematic drawing depicts the MN2a motoneuron (light green) that innervates the MHE (dark green) in an early stage 17 *Drosophila* embryo. *Right panel:* Representative confocal picture of an early stage 17 *Drosophila* embryo (*elav>GFP^RNAi^*) depicts the MN2a motoneuron (arrow) that innervates the MHE. The innervation site on the MHD muscle is highlighted by an asterisk as reference for orientation in the embryo. **(B)** Visualization of three independent genetic experiments using the pan-neural *elav*-GAL4 driver to control the expression of UAS-*RNAi* constructs of homeodomain TFs i. e. UAS-*DIPkappa^RNAi^,* UAS-*DIPgamma^RNAi^* and UAS*-GFP^RNAi^* as control. *Left panel:* Zoom on MHE of an early stage 17 *Drosophila* embryo. Note the upper panel shows a zoom of the image depicted in Figure 5A as control. *Right panel:* Schematic drawing depicts the innervation phenotype on MHE by the MN2a motoneuron. See classification of phenotypes in Figure 3B. In UAS-*DIPkappa^RNAi^,* UAS-*DIPgamma^RNAi^* animals MN2a motoneurons display axon ‘terminal defects’ and ‘wiring defects’, i.e. incorrect targeting to the LR muscle. **(C)** Representative confocal picture depicts the innervation site on the MHE muscle in L3 larvae. In animals depleted of *DIPkappa* in neurons (*elav>DIPkappa^RNAi^*) enlarged innervation sites are observed and an increased number of boutons and branching phenotypes (right panel) compared to control animals (*elav>GFP^RNAi^*) (left panel). **(D)** Representative confocal images of the pan-neural *elav*-GAL4 driver controlling the expression of the Ig domain protein dpr1 that is not expressed in Mn2a. The control experiment (*elav>GFP^RNAi^*) is depicted in Figure 5B. *Left panel:* Zoom on MHE of an early stage 17 *Drosophila* embryo. *Right panel:* Schematic drawing depicts the innervation phenotype on MHE by the MN2a motoneuron. See Figure 3B for classification of phenotypes. In wild-type conditions the MN2a motoneuron projects to the MHE, while in *elav>dpr1* animals MN2a motoneurons display ‘wiring defects’, i.e. incorrect targeting to the LR muscle. **(E)** Innervation rates for MN2a projections are calculated for *elav>DIPkappa^RNAi^* (n=17), *elav>DIPgamma^RNAi^* (n=13) and *elav>dpr1* (n=9) animals as described in Figure 3C. **(F)** Hatching rates of animals depleted of DIPkappa (*elav>DIPkappa^RNAi^*; n=165) and DIPgamma (*elav>DIPgamma^RNAi^*; n=210) in neurons compared to control animals (mock; n=211) was calculated as described in Figure 3D.

Together these results demonstrate that Ig domain proteins mediate synaptic target specificity in the neuromuscular system downstream of homeodomain TFs.

### Homeodomain TFs mediate target specificity in both matching partners

The homeodomain TFs Dfd, Mirr, and Hth are co-expressed in both functionally connected cells of the feeding motor unit, the MN2A neuron and the MHE muscle (Fig. 7A). By contrast, the adjacent LR and mouth hook depressor (MHD) muscles do not express these homeodomain TFs. This observation led us to hypothesize that combinations of homeodomain TFs label defined synaptic partners at aligned positions. To examine this hypothesis and complement the dataset on motoneurons, we performed single-cell RNA-seq of embryonic somatic muscles with enrichment for *Hox* genes, as described above for motoneurons. To this end, we sorted somatic muscles using a fly stock expressing endogenously *GFP* tagged *Myosin heavy chain* (*Mhc-TAU-GFP*, (Chen & Olson, 2001)). Analysis of this dataset by means of t-SNE and clustering indicates the existence of six relatively distinct subtypes of somatic muscle (Fig. 6A). Using previously described marker genes, these were tentatively identified as Dr+ dorsal somatic muscle, lms+ lateral somatic muscle, mid+ and well as Poxm+ ventral and lateral somatic muscle, Ptx1+ ventral somatic muscle, as well as esg+ara+ progenitor cells. For further analysis, we focused on the Poxm+ cluster and demonstrated that homeodomain TFs and Ig domain encoding genes were highly variably expressed within otherwise homogeneous somatic muscle subtypes (Fig. 6B). Most other clusters showed similar results (Fig. 6C). In the same line, we found the homeodomain TF Ubx enriched on Ig targets in the mesodermal tissue of late stage embryos in the previously generated Ubx ChIP data set (Domsch et al., 2019) (Figure 6D). These findings strongly support the hypothesis that homeodomain TF combinations are associated with *Ig* gene expression in muscles as well as in motoneurons.

**Figure 6:**
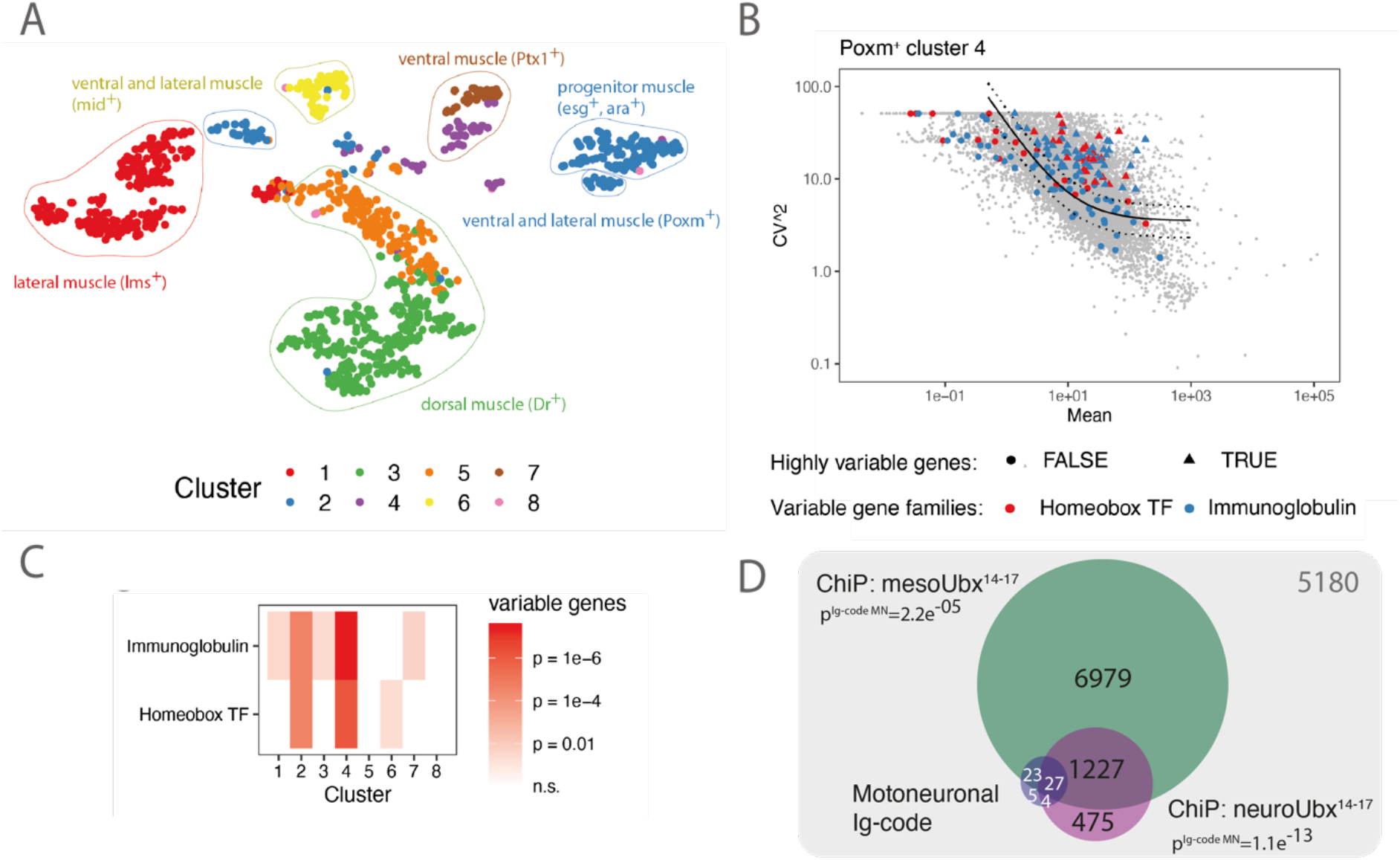
Homedomain TFs are differential expressed in the neuromuscular system and define axon target destination. **(A)** T-distributed stochastic neighbor embedding (t-SNE) plot of single-cell transcriptomes of GFP expressing somatic muscle cells sorted from a *Mhc-TAU-GFP* expressing fly line. Colors correspond to clusters identified using hierarchical clustering that were annotated using marker gene expression of muscle subtypes. **(B)** Statistical model of Brennecke et al. (2013) was used to identify highly variable genes (see panel D for an example). Color code displays enrichment of Immunoglobulins or homeobox TFs among variable genes according to a hypergeometric test. **(C)** Identification of highly variable genes using the method by (Brennecke et al., 2013). Scatter plot depicts for each gene the mean expression and squared coefficient of variation across cells from the Poxm+ cluster 4. The solid line indicates the fit, dashed lines the 95% confidence interval. Genes with a significantly elevated variance are shown as triangles, other genes as circles. Different gene classes are color coded. **(D)** Venn diagram displays the enrichment of Ig domain genes bound by Ubx in a muscle and neuronal tissue specific ChIP (Domsch et al., 2018), n=13920 genes (all protein coding genes expressed in the *Drosophila* genome).

**Figure 7:**
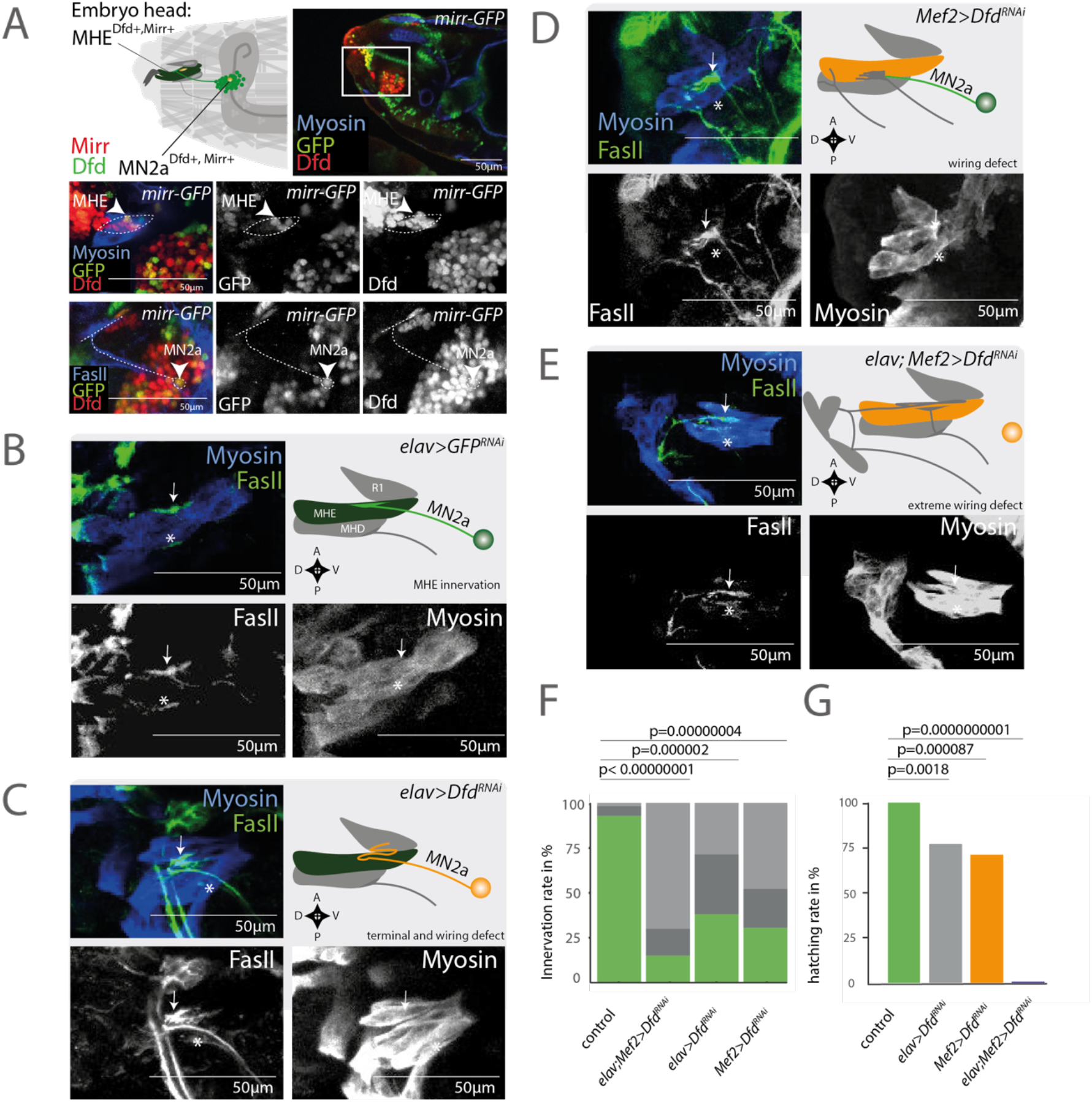
Synaptic targeting requires the homeodomain TF Dfd in neurons and muscles. **(A)** *Left panel:* Schematic drawing depicts the MN2a motoneuron (yellow) expressing the homeodomain TFs Dfd (green) and Mirr (red) that innervates the MHE (yellow nucleus) expressing the same set of homeodomain TFs in an early stage 17 *Drosophila* embryo. *Right panel:* Representative confocal picture of an early stage 17 *Drosophila* embryo (*mirr-GFP*). The mid panel and lower panel show two independent experiments i. e. mid panel displays expression of the homeodomain TFs Mirr and Dfd in the MHE (arrow), while the lower panel depicts expression of Mirr and Dfd in the MN2a motoneuron that innervates the MHE. The MHD muscle underneath the MHE is negative for Mirr and Dfd**. (B-E)** Visualization of three independent genetic experiments using the pan-neural *elav*-GAL4 driver, the muscle driver (*Mef2-*GAL4) and both of these drivers combined (*elav-*GAL4*; Mef2-*GAL4) to control the expression of a UAS-*Dfd^RNAi^* construct. *Left panel:* Zoom on MHE of an early stage 17 *Drosophila* embryo. Note panel 6b and 6c show the same images as depicted in Figure 5B. *Right panel:* Schematic drawing depicts the innervation phenotype on MHE by the MN2a motoneuron. See classification of phenotypes in Figure 3B. In *elav>Dfd^RNAi^* animals MN2a motoneurons display axon terminal defects and ectopic targeting to the LR muscle, while *Mef2>Dfd^RNAi^* and *elav;Mef2>Dfd^RNAi^* show extreme wiring defects and ectopic innervations on the MHE muscle target. **(F)** Innervation rates for MN2a projections are calculated for *elav>Dfd^RNAi^* (n=21), *Mef2>Dfd^RNAi^* (n=23) and *elav;Mef2>Dfd^RNAi^* (n=20) animals as described in Figure 3C. **(G)** Hatching rates of animals depleted of Dfd in neurons (*elav>Dfd^RNAi^;* n=217), in muscle (*Mef2>Dfd^RNAi^*; n=284) in neurons and in muscle (*elav;Mef2>Dfd^RNAi^*; n=2403) compared to control animals (mock; n=2898) was calculated as described in Figure 3D.

To further study the role of homeodomain TFs in both muscles and neurons, we interfered with the expression of *Dfd* in a tissue dependent manner to examine synaptic defects of stage 17 embryos. Hence, we knocked down *Dfd* (UAS-*Dfd^RNAi^*) individually in neurons (*elav*-GAL4) and in muscles (*Mef2*-GAL4) or in both tissues (*elav*-GAL4; *Mef2*-GAL4;). In all three conditions we observed significant synaptic wiring defects and terminal defects (Fig. 7B-7F). Interestingly, the strength and type of wiring defects differed in each tissue. *Dfd* depletion in neurons and muscles alone showed a wiring defect in 29% and 48% of the embryos respectively, while knockdown of *Dfd* in both tissues increased the rate to 70% of cases, indicating that homeodomain TFs are required in both tissues neurons and muscle to coordinate synaptic partner matching. *Dfd* depletion in MN2a led to mistargeting of the LR muscle (Fig. 7C), while *Dfd* depletion in muscles led to innervation of the MHE muscles by motoneurons that were supposed to innervate the MHD muscles (Fig. 7D). Knockdown of *Dfd* in both muscle and neurons caused aberrant innervations of the MHE muscles and loss of the stereotypic innervation (Fig. 7E), suggesting that homeodomain TF expression in both tissues is required to regulate repellent and adhesive factors for synaptic target selection thereby preventing mistargeting. In all three conditions, hatching rates were significantly decreased: by 23% when *Dfd* was depleted in neurons, by 29% when *Dfd* levels were reduced in muscle, and by 99% when *Dfd* was silenced in both tissues (Fig. 7G). These results demonstrated the importance of homeodomain TF expression in both neurons and muscles for synaptic wiring and behavior.

In sum, the results showed that combinations of homeodomain TFs are required to align synaptic partners in the neuronal and muscle tissues and coordinate their preferences in synaptic partner choice.

## DISCUSSION

Each neuron in the nervous system chooses a single target cell from a high number of possible interactions, which is a prerequisite for the formation of stereotypic neuronal circuits. This extraordinary degree of precision is thought to be mediated by a matching code of cell recognition and adhesion molecules such as CSPs. However, up to date a systematic scrutinization of this code and upstream mechanisms fine-tuning its expression are missing. The most important reasons for this are the lack of single cell specific neuronal markers, the complexity of neuronal systems and the possibly gradual and combinatorial nature of CSP expression.

Our experimental design allowed us to overcome these challenges. First, by focusing on one neuronal subtype, the motoneuronal population in *Drosophila* embryos, we were able to reduce neuronal complexity. Second, we investigated this cell population exactly at the time when they form stereotypic connections with their muscle targets, which allowed us to identify molecular cues critical for synaptic wiring. Third, motoneurons form highly cell-specific connections with muscles and are present in a relatively small number per embryo. Thus, we used a single-cell genomic approach with a high number of biological replicates of every biologically unique cell. Thereby we identified novel markers specific to single cells or small groups of cells that in turn permit the identification of transcriptome signatures relevant for wiring. And finally, we implemented a spatial mapping approach based on Hox gene expression to locate motoneurons along their AP position and thereby gained insight into the role of spatial mechanisms during synaptic wiring.

Our scRNA-Seq data revealed that a homeo-code acts as main determinant for transcriptional heterogeneity during the wiring phase of motoneurons. scRNA-Seq, despite the use of a high number of biological replicates for each unique cell, cannot be applied to unanimously identify biologically unique cells. Thus, we used imaging to demonstrate its specificity to single cells in six out of six cases investigated. Together, these data allowed us to conclude that a homeo-code defines cellular identities in *Drosophila* motoneurons, possible down to the level of single cells. Recent work has demonstrated the existence of a similar code in *C. elegans*, suggesting that homeodomain TFs might be utilized to specify unique neuronal cell identities throughout evolution. Indeed, homeodomain TFs are known to play a role in the specification of mammalian neuronal subtypes as well (Buelow et al., 2005; Dasen, 2018; Mallo, 2014; Song & Pfaff, 2005).

To gain insight into the developmental origin of the homeo-code, we spatially mapped all motoneurons along the AP axis using *Hox* gene expression as markers. We validated the high accuracy of our mapping approach, and point out that our strategy requires a much smaller number of markers compared to existing approaches for the spatial mapping of single-cell transcriptomes (Achim et al., 2015; Bageritz et al., 2019; Satija et al., 2015). We found that position along the AP axis decisively impacts the expression of homeodomain TFs at the synaptic wiring stage. Furthermore, genes associated with the dorsal-ventral position of motoneurons are highly variable but statistically independent of AP position. These results suggest that during development, molecular position is imprinted early on and affects cellular function during synaptic wiring. However, we also observed substantial additional variability of homeodomain TF expression, suggesting that beyond such spatial mechanisms, other processes induce the specification of unique cellular identities, which could be neuronal birth order.

Homeodomain TF expression is not only highly specific to unique cellular identities, but is also associated with profound differences in the entire transcriptome and in particular the expression of Ig CSPs as possible effectors of synaptic specificity. Importantly, manipulation of homeodomain TF expression during the wiring phase by knockdown or ectopic expression leads to changes in target preferences, which are phenocopied by manipulations of their Ig targets. Together, the combination of single cell genomics (co-expression), chromatin immunoprecipitation (binding of enhancers) and genetics (phenocopies) demonstrate that while the homeo-code specifies unique cellular identities, Ig proteins act as the effectors of synaptic specificity.

Classification of cellular identities using the homeo-code allowed us to systematically investigate the CSP distribution between cells. Interestingly, most CSPs change more gradually between cells, and unlike homeodomain TF expression, the binary expression of Ig proteins alone is insufficient to specify single cells (De Wit & Ghosh, 2016; van Oostrum et al., 2020). Previous studies on single molecules even indicated that already small changes in relative expression levels of CSPs in matching partners can change synaptic specificity (Sweeney et al., 2012; Yogev & Shen, 2014). Here only one Ig molecule class, the DIP genes, were found to be distinct for specific cellular identities and genetic manipulations on single candidates are required to modulate synaptic specificity and synaptic affinity to target cells (ON-Mode). Furthermore, we found in our data set some Ig genes, such as Dscam encoding genes, to be broadly expressed but not present in some defined cells (OFF-Mode). Finally, most other Ig genes are less specific for cellular identities and relative expression levels change gradually between cells, as suggested previously (De Wit & Ghosh, 2016; van Oostrum et al., 2020). In sum, our data show that a highly combinatorial Ig code drives synaptic specificity and connectivity between cells.

Our data justify a model whereby the position of every cell is imprinted early in embryonic development by patterns of homeodomain TFs (Fig. 8). Expression patterns of these factors become more complex and combinatorial with each cell division until small groups of cells and possibly every single cell is uniquely labelled by a homeo-code. These unique combinations of homeodomain TFs then in turn regulate specific downstream programs of immunoglobulin gene expression. We have shown that in the target cells (here: muscles), connectivity is functionally specified by a similar transcription factor code, which possibly induces the expression of a complementary Ig receptor expression program. This molecular logic enables every single cell to find its corresponding interaction partner based on complementary adhesive properties mediated by combinations of Ig domain molecules. In sum, this concept explains how a molecular memory of cell body position is translated into invariant cell-cell adhesion events by means of the homeo-Ig-code.

**Figure 8:**
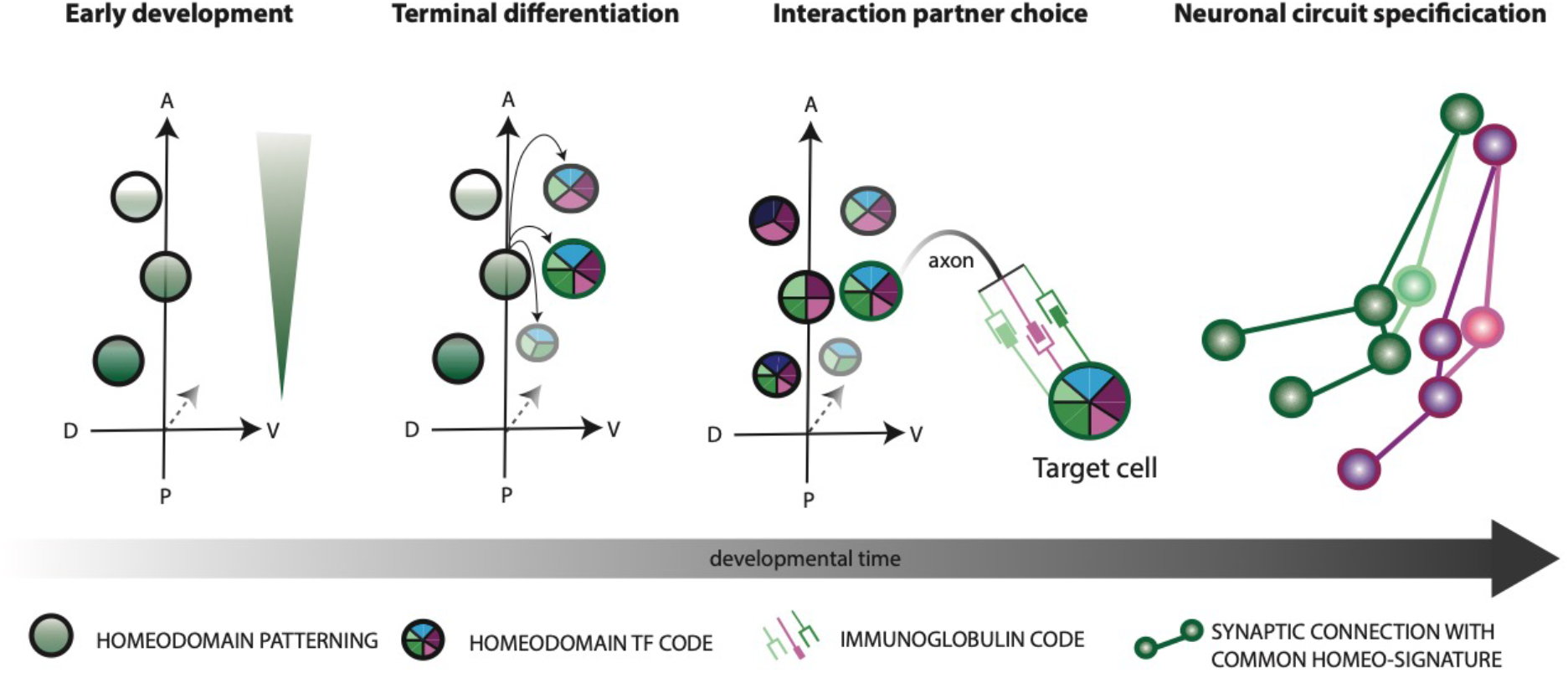
The molecular mechanism for stereotyped synaptic partner selection and neuro-muscular circuit architecture. *Left panel:* Early developmental programs pattern single cells according to their position in the embryo. Morphogenic gradients establish specific homeodomain patterns along embryonic body axis (i.e. dorsal ventral axis = DV axis, anteroposterior axis = AP axis). *Left middle panel:* After several divisions of neuroblasts, neurons terminally differentiate and establish unique identities with distinct homeodomain TF codes (color code). *Right middle panel:* A unique homeodomain TF code specifies Ig domain receptor expression in matching synaptic partners and differential affinities of Ig domain proteins promote selective synaptic target choice. *Right panel:* Common homeo-signatures specify interaction partners within individual neuro-muscular circuits.

Our data suggest that the structure of neuronal circuit is encoded in the gene regulatory logic of Ig domain encoding genes. How their regulation by homeo-TFs is hardwired into their cis -or trans-regulatory elements is a topic for future investigation. In particular, it is unclear how specificity is achieved, despite homeodomain TFs binding to relatively similar motifs. In the future, the combinatorial manipulation of CSPs and homeodomain TFs in single cells (Replogle et al., 2020) will allow a further functional dissection of their interactions and regulatory mechanisms.

## ACKNOWLEDGMENTS

We thank V. Benes and his team at EMBL Gene Core for performing the next generation single RNA sequencing for this study. We thank D. Ordones at EMBL for running the FACS sorting experiments of the muscle data set. We thank the Proteomics Facility at EMBL for providing the homemade Tn5 enzyme for single cell library preparation. We thank S. Sorge for cloning the TRiP-vector for the Dfd^RNAi^ fly line. We thank T. Schunke for providing the initial version of the single cell dissociation protocol. We thank P. Kaspar for ordering and managing all the equipment and B. Glaß for cooking the fly food required for this study. We thank J. Friedrich and S. Sorge for the initial introduction into the topic. We also thank A. Teleman and S. Lemke for critically reading the manuscript. The project was funded by the DFG grant LO 844/4-2 to Ingrid Lohmann.

## AUTHOR CONTRIBUTIONS

J.V. and I.L. designed the study with contributions by L.V.. J.V. performed experiments with contributions by R.A., L.B. and M.P.. J.V. and L.V. performed data analysis. P.V.N. contributed code for image analysis. J.V. wrote the paper together with I.L. and L.V.. I.L. obtained funding for this study (DFG, LO 844/4-2). All authors have read and commented on the paper.

## METHODS

### *Drosophila* strains and experimental crosses

We used the *OK371*-GAL4 driver (Mahr & Aberle, 2006) crossed to UAS-*mCD8-RFP* to perform single cell sorting of motoneurons, and the *Mhc-TAU-GFP* line (Chen & Olson, 2001) for sorting of somatic muscle cells. Crosses were kept for 1 hours at 25°C to oviposit on apple-juice plates with yeast paste. Subsequently eggs were incubated on apple-juice plates for further 19 hours and then embryos were dissociated for FACS sorting.

For genetic experiments, we used the pan-neuronal *elav-*GAL4 driver (Luo et al., 1994) and the muscle specific *Mef2-*GAL4 driver (Ranganayakulu et al., 1998) as main driver. To confirm phenotypes in motoneurons the *OK6-*GAL4 driver (Mahr & Aberle, 2006) was used in a few examples. In this study we used second generation *TRiP RNAi* lines from Bloomington (Ni et al., 2011), because these *short hairpin RNA* (*shRNA*) are more effective in embryonic stages than long hairpin of first generation of the TRiP project based on own experiences. Therefore, only one UAS-*RNAi* line fulfilling the required criteria has been available for most of the target genes. The UAS-*Ubx-HA* line is described in (Domsch et al., 2019). For innervation rate assays we crossed virgins of the corresponding driver line to males carrying UAS-*RNAi* or overexpression constructs. As Control we performed the experiment under the same conditions with UAS-*mcd8-RFP* lines or UAS-*EGFP^RNAi^* lines. The crosses were kept for at least 16 hours at 29°C for knockdown experiments and 20 hours at 25°C for overexpression experiments on apple-juice plates. Early stage 17 embryos were selected after embryo fixation.

For hatching rate assays, the *elav*-GAL4 driver, *Mef2*-GAL4 and the *elav*-GAL4; *Mef2*-GAL4 driver (designed for this study) were crossed to UAS-*Dfd^RNAi^* (designed using the second generation of TRiP system) or to Bloomington lines UAS-*DIPgamma^RNAi^* and UAS-*DIPkappa^RNAi^*. UAS-*mcd8-RFP* lines or UAS-*EGFP^RNAi^* lines crossed to the above-mentioned driver lines were used as controls. Crosses were set up in duplicates (equal amounts of males and females for every replicate, sample and control, 1:1 ratio) and egg laying was performed at 29°C for 3 hours on apple-juice plates with yeast paste. Subsequently, eggs were washed, counted and unfertilized eggs were removed or counted and transferred on fresh apple juice plates without yeast paste. Eggs were incubated for additional 24 hours and then the hatching rate was quantified.

For characterization of synaptic phenotypes in larvae (L3), we crossed *elav*-GAL4 to UAS-*Dfd^RNAi^* or UAS-*DIPkappa^RNAi^* and developed progeny for about 6 days at 25°C for larval dissections and fixation.

### Plasmid construction and transgenesis

The UAS-*Dfd^RNAi^* line was generated following the conventional TRiP protocol (second generation).

### Immunohistochemistry

Immunofluorescence experiments on *Drosophila* late stage embryos and mouth hook muscles and central nervous system of late L3 larvae was performed according to standard protocols.

The embryo fixation protocol in brief. First embryos were bleached for 2 min to remove the chorion. After washing in water, embryos were transferred to fixing solution (3,7% Formaldehyde in PBS + 100% Heptane) and incubated for 20 min at room temperature on a nutator. Fixing is stopped by removal of Formaldehyde. Then equal amounts of Methanol are added to Heptane and vortexed for about 40 seconds to remove the vitellin membrane. Subsequently the Heptane phase is removed, and embryos are washed in Methanol.

For antibody stainings, embryos were hydrated and washed for 3 times in PBT (with 0.1% Tween 20). The primary antibodies were used at 4°C overnight from Abcam (rat anti-Myosin 1:1000) DSHB (mouse anti-FasII, 1:50; mouse anti-Scr, 1:50; mouse anti-Abd A, 1:50; mouse anti-Antp; rat anti-Elav 1:50), Invitrogen (rabbit anti-GFP, 1:500), produced by Katrin Domsch (guinea pig anti-Dfd 1:500; guinea pig anti-Ubx 1:500; anti-Lab 1:500) or unknown source (anti-Abd B, anti-Hth, rat anti-Vvl). After 3x washing in PBT (with 0.1% Tween 20) embryos were incubated for 2 hours at room temperature with secondary antibodies from Jackson Immunoresearch. Vectashield with DAPI or TSO was used as mounting medium.

For double stainings with antibodies originating from the same animal (mouse anti-FasII + mouse anti-Antp; mouse anti-FasII + mouse anti-Abd A; mouse anti-FasII + mouse anti-Scr), we performed a modified protocol for sequential antibody staining by TSA (according to the manufacture protocol) The central nervous system and mouth hooks with mouth hook muscles attached were dissected from L3 larvae. The mouth hooks with larval brains were fixed with 4% paraformaldehyde for 20 min at room temperature and subsequently washed 3x with PBT (0.3% Triton-100). Primary and secondary antibodies were used with similar concentrations than for antibody staining’s of embryos.

### Microscopy and image analysis

Fixed embryos and larval brains with mouth hook muscles were imaged by Leica TSC SP8 confocal microscope. The 20x Objective was used for imaging of embryos and the 63x objective to visualize the neuromuscular junction of the mouth hook elevator muscle. Images were processed by Fiji. Embryos used for quantification of innervation rate assay were processed with a standardized imaging pipeline (programed by Patrick van Nierop). Confocal pictures shown in figures were processed manually to optimize the result.

For AP axis measurements along the ventral nerve cord, we used FasII staining as reference and measure protein intensities of Lab, Dfd, Scr, Antp, Ubx, Abd A and Abd B by Fiji. We normalized the length of the ventral nerve cord between different embryos (0=anterior to 2000=posterior), set thresholds for the protein intensities to remove the overall unspecific background caused by antibody staining’s and normalized the protein intensities of different source antibodies (0=min intensity to 100=max intensity).

### Dissociation of embryonic cells and Flow cytometry

Collections of 19-20h old *Drosophila* embryos were dechorionated with bleach (5%Chloride) for 2 min. For dissociation embryos were transferred into syringes filled with 1ml of 10x Trypsin-EDTA (0,5% Trypsin). For additional mechanical dissociation embryos were transferred 20x between two syringes (25G needle) and 10x between two syringes with 27G needles. Debris was reduced by filtering dissociated cell via cell strainer. The cells were stained for 5 min with DRAQ5 (1mg/ml) and DAPI (1mg/ml). Subsequently cells were sorted into 96-well plates containing 5 µl of Smart Seq2 lysis buffer at 4°C by a BD FACS Aria III flow cytometer equipped with 405nm, 488nm, 561nm and 633nm laser. Directly after cell sorting plates were shock frozen in nitrogen.

### Single-cell transcriptome sequencing

A pooled cell population of Ok371>RFP positive motoneuronal cells and a pooled population of Mhc-TAU-GFP positive somatic muscle cells derived from FACS-sorted 19-20h old *Drosophila* embryos was used for scRNA Sequencing. The standard smart-seq2 protocol (Picelli et al., 2014) was modified by targeting low expressed Hox genes during the reverse transcription and PCR stages (*lab, Dfd, Scr, Antp, Ubx, abdA, AbdB*). Therefore, 1µM targeted primer mix of each *Hox gene* (See key resource table) was added to the PCR and RT buffers, respectively. The above described approach targets all expressed isoforms of each Hox gene. Method specific biases were ruled out by quality checks via Bioanalyzer of the whole transcriptome after SmartSeq2 preparation. Libraries of scRNA transcriptomes were created using a home-made Tn5 transposase (Hennig et al., 2018) and seventeen 96 well plates were sequenced with 75bp single-end on an Illumina NextSeq platform.

### Raw data processing, quality control and normalization

Sequencing reads were demultiplexed and the Poly-A tail trimmed. Read count tables were generated by pseudo-alignment to the cDNA of the *Drosophila* transcriptome (BDGP6 ensemble) using Kallisto (Bray et al., 2016). For quality control, we used standard settings (Velten et al., 2017): Cell were removed unless they contained at least 10 reads for each of at least 500 genes, and genes were removed unless they were expressed in at least 5 cells with 10 reads each. Data was normalized and scaled using the indexplorer pipeline (Velten et al., 2017). Posterior odds ratio scaling (Velten et al., 2017) was used to scale variance according to an estimate of true biological variance, as opposed to technical variance.

### Clustering and dimensionality reduction

Following the indexplorer pipeline, PCA was computed using scaled, normalized expression values of all genes. t-SNE was then computed on the first ten principal components based on a visual inspection of the PCA Elbow plot. Alternatively, and following (Li et al., 2017), cells were clustered by hierarchical clustering using only the 20 most variably expressed genes of our scRNA data set; Ward linkage was used on an Euclidean distance metric.

### Inference of spatial position from single cell gene expression data

To infer spatial position from single cell gene expression data, we first created a reference map of protein expression for seven *Hox* genes (Supplementary Figs 3a, 3b). To that end, immunofluorescence experiments were performed as described above.

Immunofluorescence data for gene *g* was thereby represented as a function of position *Y*_g_(*x*) ∈ (0,1) whereas single cell gene expression was represented as a matrix of read count values across genes and cells, *D_g,c_* ∈ ℕ_0_. We then assume that the probability of observing *D_g,c_* given that the position of cell *c* is really *x_c_* depends on the expression of *g* at that position:

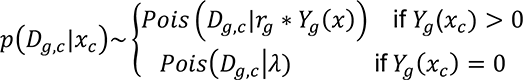

Here, *λ* is a constant corresponding to a background number of reads observed in non-expressing cells, and *r_g_* is a gene-wise scaling factor that estimates the average number of RNA-seq reads in a cell maximally expressing the protein. A maximum likelihood estimate of *x_c_* is then obtained by computing

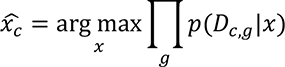

For the analysis shown in the manuscript, *λ* was set to 0.1 (Kharchenko et al., 2014) and *r_g_* was set to the mean gene expression in expressing cells:

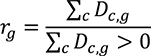

We used Latin Hypercube Sampling across a wide range of values for *λ* and *r* to demonstrate that the position estimate was independent of parameter choice (not shown).

To identify genes with spatially variable expression, B-Spline models with 3 degrees of freedom were fitted through scaled and normalized gene expression values for each gene individually, using inferred position as the only covariate. These models were then compared to null models not containing the position term using the Bayesian Information Criterion, similar to the workflow for selecting genes with variable expression over pseudotime described in (Velten et al., 2017). For visualization and clustering, expression values for all variably expressed genes were arranged by inferred position and a floating mean was computed by 1D-convolution with an absolute exponential kernel with decay rate 10. Smoothened gene expression values obtained thereby were then compared to immunofluorescence images for model validation (Fig. 1E) or used for clustering of gene into modules with coherent expression patterns over space (Fig. 1f).

### ZiNB-WAVE analysis

To identify gene whose expression was variable but statistically independent from the AP axis, we made use of the ZiNB-WAVE model (Risso et al., 2019). Unlike a PCA, ZiNB-WAVE separately estimates a matrix of known-covariate factors as well as a matrix of unknown-covariate factors; furthermore, ZiNB-WAVE uses a zero-infalted negative binomial distribution to account for the sparse nature of single cell RNA-seq data. We ran ZiNB-WAVE using default settings and the number of genes observed as well as the inferred AP position as known sample-level covariates. While the PCA of the dataset was dominated by effects related to the number of genes observed, viability dye incorporation, and AP position (see Supplementary Fig. 1), the unknown-covariate factors from ZiNB-WAVE arranged genes according to their dorsal-ventral expression.

### ChIP-Seq reanalysis and iRegulon analysis

To identify the reletionship between motoneuronal genes expressed in our scRNA-Seq data set and genes bound by the homeodomain TF Dfd, we used a whole embryo Dfd ChIP (Sorge et al., 2012) to investigate if Ig domain proteins are overrepresented among putative Dfd-regulated genes compared to other genes expressed in motoneurons. Genes were classified as regulated by Dfd, if Dfd is bound to the target gene, max. 1kb upstream of the promotor region (Sorge et al., 2015). To calculate a p-value for enrichment, we performed a hypergeometric test.

The iRegulon (Janky et al., 2014) analysis was used to identify TF motifs enriched in the vicinity of Ig encoding genes expressed in our motoneuronal scRNA data set. Therefore, we performed an iRegulon analysis on all Ig domain encoding proteins expressed in motoneurons under standard settings (9,713PWM; 5kb upstream, 50 UTR and first intron with standard cutoff) to identify the 15 highest ranked transcription factor motifs. Then we used iRegulon to investigate which TF families are predicted to bind to these motifs and chose 3 of the 5 motifs predicted to be regulated by homeodomain TFs.

### Data Visualization

All plots were generated using the ggplot2 (v. 3.2.1) and pheatmap (v. 1.0.12) packages in R 3.6.2. Boxplots are defined as follows: The middle line corresponds to the median; lower and upper hinges correspond to first and third quartiles. The upper whisker extends from the hinge to the largest value no further than 1.5 * IQR from the hinge (where IQR is the inter-quartile range, or distance between the first and third quartiles). The lower whisker extends from the hinge to the smallest value at most 1.5 * IQR of the hinge. Data beyond the end of the whiskers are called “outlying” points and are plotted individually.

### Code availability

Most analyses were done using the indexplorer software for interactive exploration of single cell RNA-seq datasets (Velten et al, 2017), which is available from https://git.embl.de/velten/indeXplorer. Custom scripts for inference of spatial position are available from the corresponding authors upon request.

### Data availability

RNA-seq data is available for interactive exploration at https://ie-collaborators.shiny.embl.de/?who=allNeurons (for the neuronal dataset) or https://ie-collaborators.shiny.embl.de/?who=allMuscle (for the muscle dataset). Raw data and count tables are available from gene expression omnibus, ID: GSE155578 (for the motoneuronal data set) and ID: GSE155586 (for the muscle data set) reviewer access for both data sets: svgvmiqijvobfwz.

## SUPPLEMENTARY MATERIAL

**Supplementary Figure 1:**
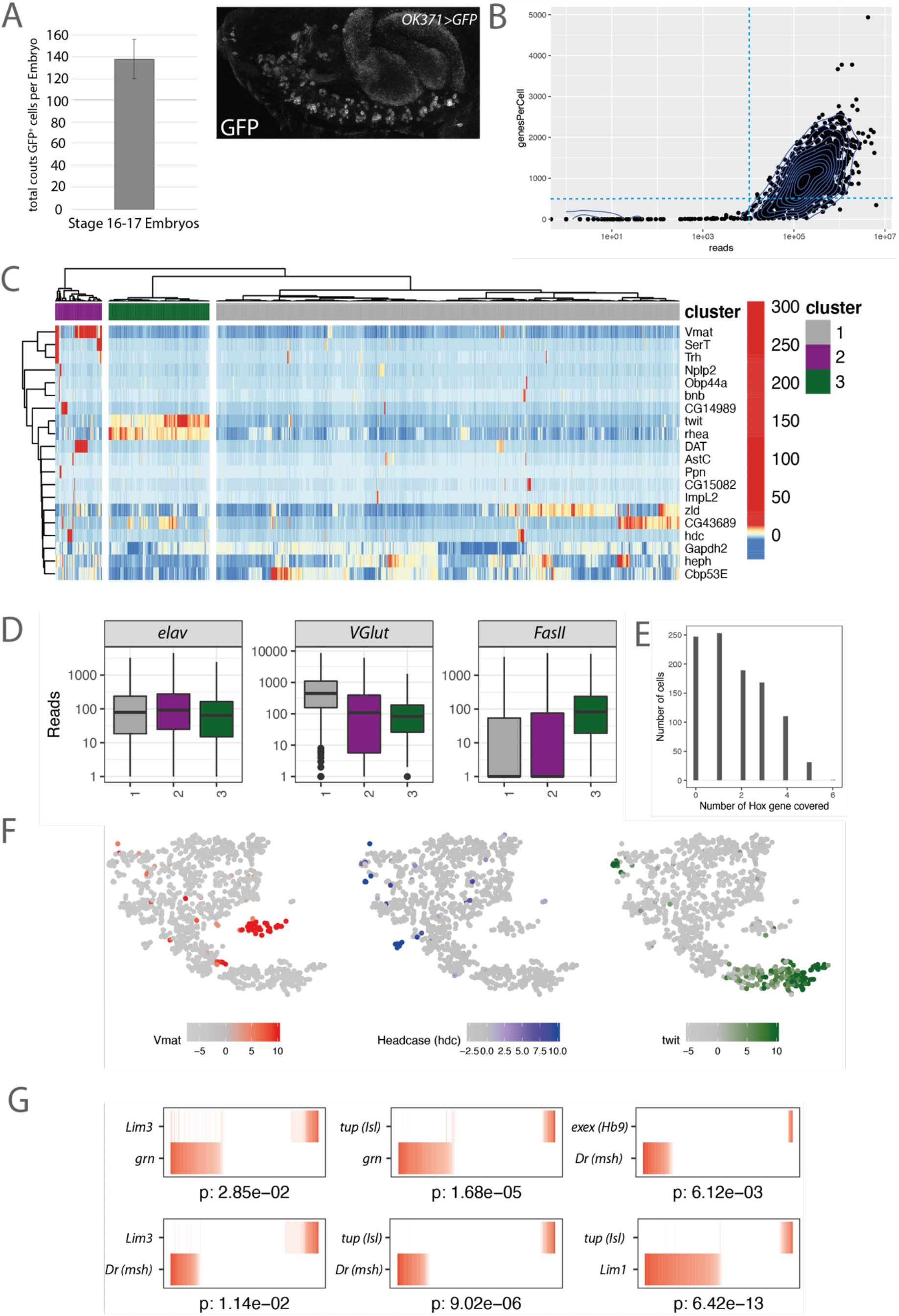
High quality single-cell RNA sequencing of embryonic *Drosophila* motoneurons. **(A)** Visualization of filtering criteria for single cells (dashed blue line, see Materials and Methods). Density dot plot represents the total reads (library size) versus genes observed per cell (library quality, diversity). Each dot represents a motoneuronal cell (total of 1536 cells). In total, 999 cells passed the filtering criteria indicated by the dotted lines (see Methods). **(B)** Quantification of the average number of terminally differentiated *OK371>GFP* positive motoneurons in stage 16-17 embryos (n=3). **(C)** Single motoneuronal cells (columns) are hierarchically clustered using the 20 most variable expressed genes (rows) following methodology of Li et al. (2017). Hierarchical clustering was performed using ward linkage on an euclidean distance metrics. Color code represents gene expression levels (see Methods). **(D)** The marker genes *elav* (pan-neural), *VGlut* (glutamatergic motoneuronal) and *FasII* (motoneuronal) were evaluated for each of the three clusters described in Figure S1C. **(E)** Bar chart represents the number of cells expressing *Hox* genes. 749 cells of 999 cells express at least one *Hox* gene (∼75% *Hox* gene coverage). **(F)** Expression level of key marker genes are highlighted on the t-SNE of Figure 1B. **(G)** Expression of gene pairs described in literature to be expressed in a mutually exclusive pattern. Columns correspond to single cells.

**Supplementary Figure 2:**
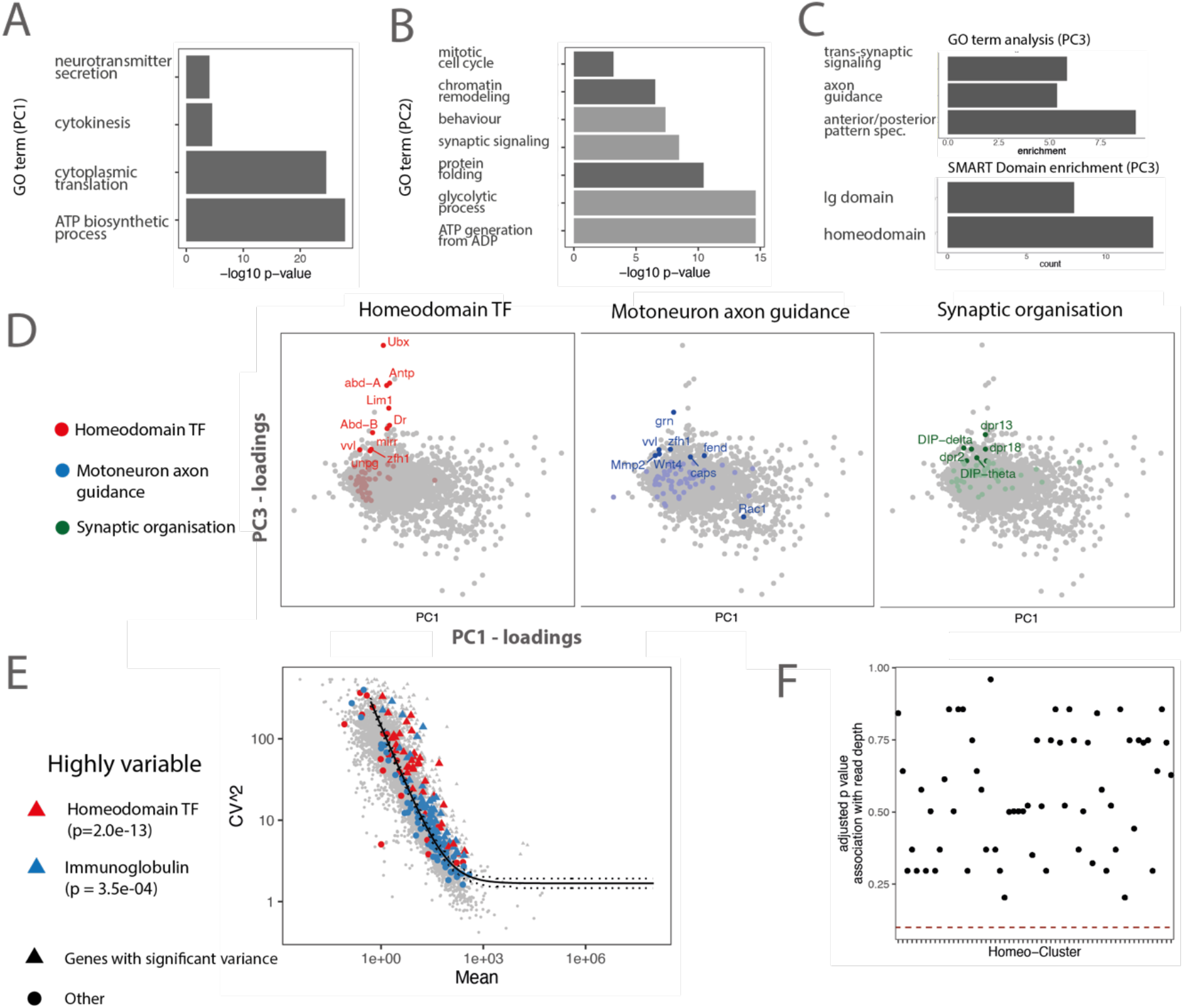
Identification of variable processes in embryonic motoneurons by principal component analysis. **(A, B, C)** Principal component analysis (PCA) was performed on cells from the *twit^low^* cluster. GO term analysis for biological processes was performed on the top 10% genes with highest loadings on principal component 1, PC1 (log 10 p-value) (A), principal component 2, PC2 (B) and principal component 3, PC3 (C). GO term and SMART domain analysis was performed on the top 300 genes representing the most enriched candidates in the PCA. Dark grey indicates processes enriched among genes with positive loadings, light grey indicates processes enriched among genes with negative loadings. Together these analyses indicated that PC1 and PC2 are associated with metabolic processes, cellular differentiation and/or technical variation, while PC3 is associated with anterior, posterior patterning processes and synaptic specificity processes. **(D)** Principal component loadings plots highlighting homeodomain TFs (red), genes associated with the GO terms motoneuron axon guidance (blue) and synaptic organisation (green). Points with label correspond to the highest 5% of loadings. **(E)** Identification of highly variable genes using the method by (Brennecke et al., 2013). Scatter plot depicts for each gene the mean expression and squared coefficient of variation across twil^ow^ cells. The solid line indicates the fit, dashed lines the 95% confidence interval. Genes with a significantly elevated variance are shown as triangles, other genes as circles. Different gene classes are color coded. P values shown are from a hypergeometric test for enrichment of the respective gene class among highly variable genes. **(F)** Scatter plot depicting for each homeo-cluster from main Figure 1C the strength of association with a technical covariate (sequencing depth). P values are from a wilcoxon test contrasting sequencing depth in cells from that cluster, and all other cells. The dotted read line indicates the p value required for significance (0.05). All associations are therefore non-significant.

**Supplementary Figure 3:**
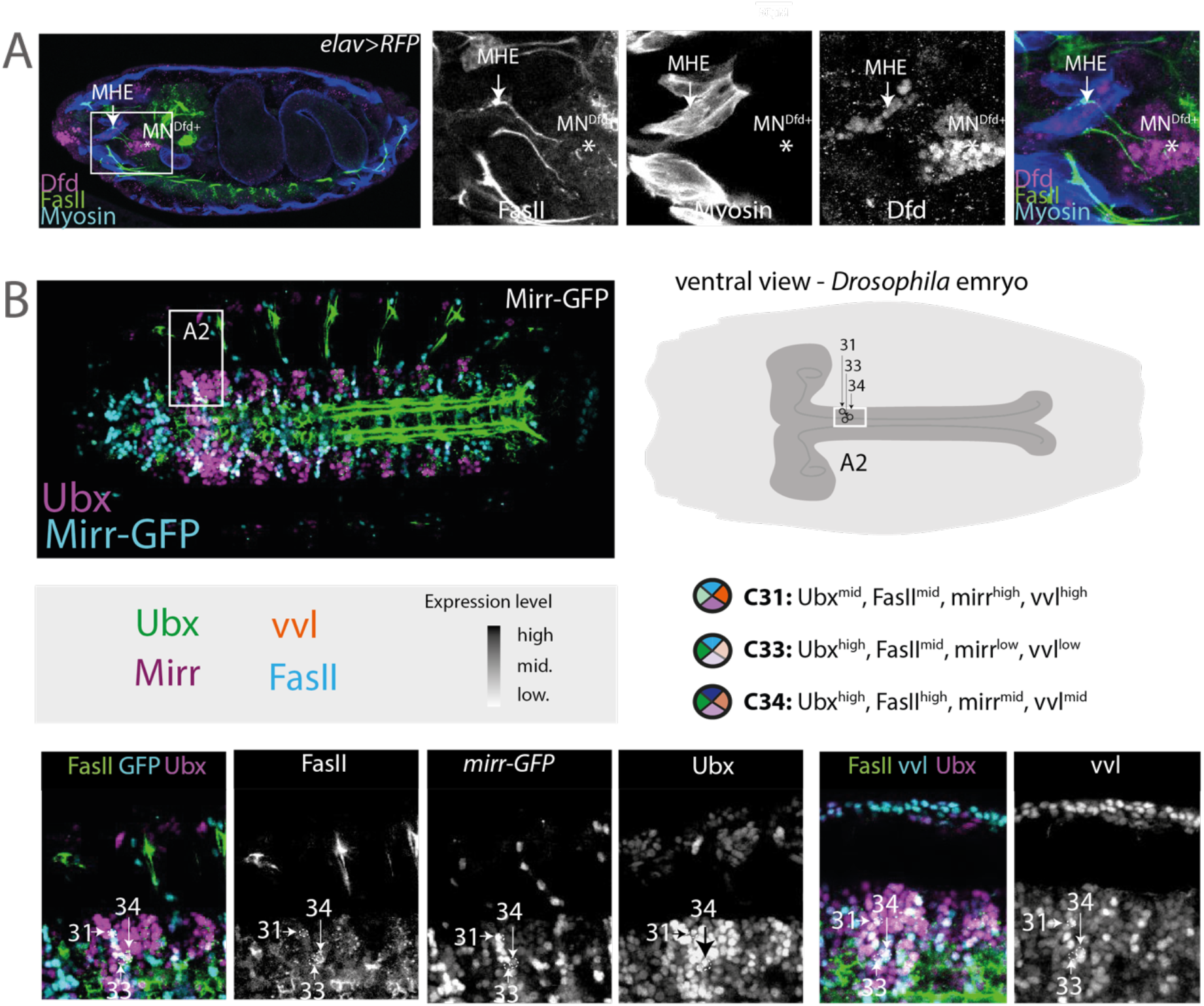
Validation of homeo-cluster in single cells. **(A)** Confocal image analysis of the lateral view of early stage 17 *Drosophila* embryos to visualize Dfd expression (magenta) in FasII-positive motoneurons (green) and Myosin-positive muscles (blue) for the identification of the exact target destination of the Dfd^+^ MN2a motoneuron. The MN2a axon (asterisk, Figure 2F) projects to the MHE (arrow). **(B)** *Left upper panel:* Confocal image depict the ventral view of early stage 17 *Drosophila* embryos. Motoneurons are labelled by FasII (green). *Right upper panel:* Schematic drawing and immunofluorescence image depict the ventral view on an early stage 17 *Drosophila* embryo and highlights the A2 segment in the ventral nerve cord and the location of the three validated cells below. *Lower panel:* Zooms on the abdominal A2 segments and highlights the combinatorial expression of homeodomain TFs (Mirr, Vvl and Ubx) expression in individual cells (white arrows) that corresponds to expression levels of clusters 31, 33 and 34 of scRNA data depicted in Figure 1C.

**Supplementary Figure 4:**
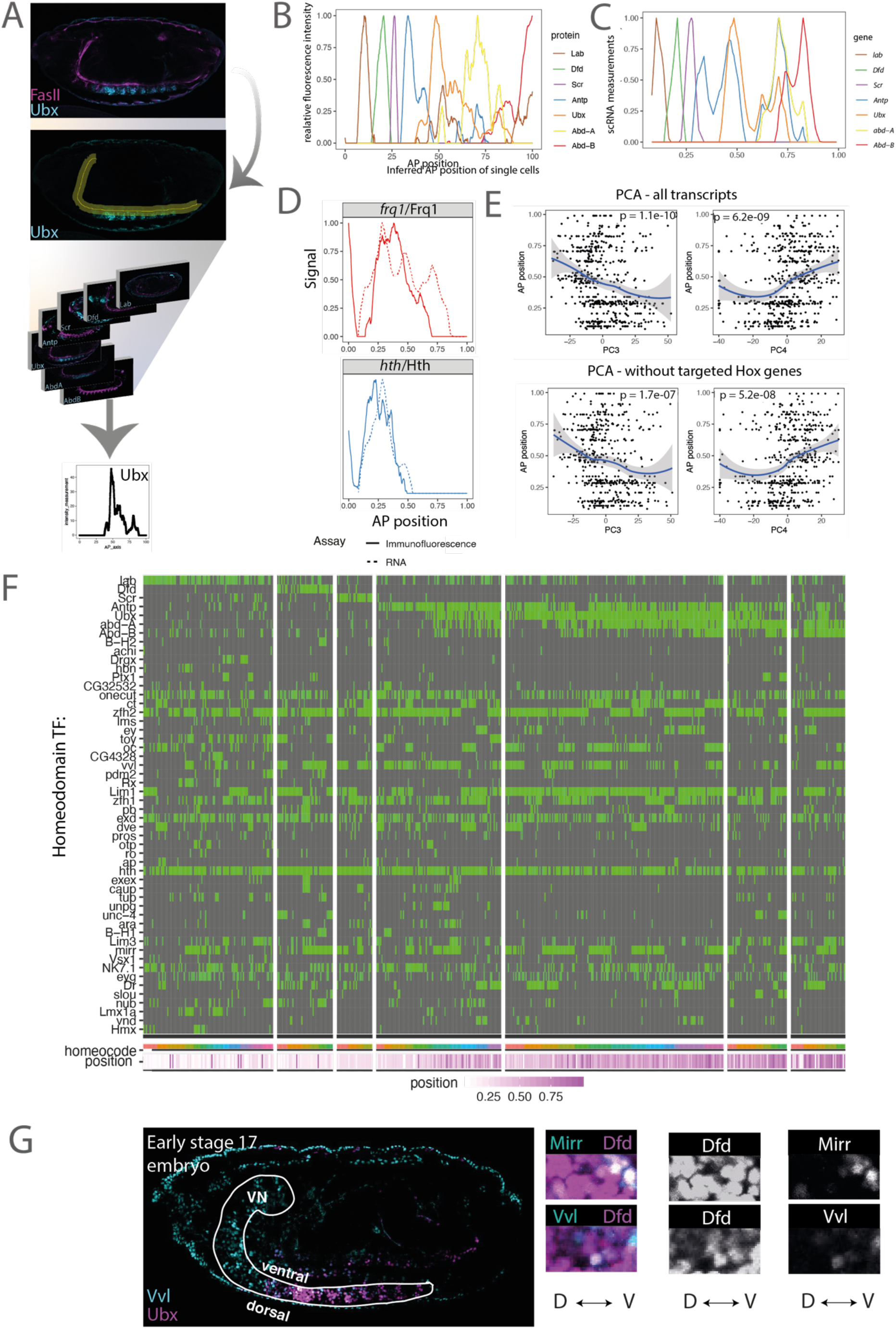
Spatial reconstruction of AP axis based on Immunofluorescence measurements of Hox proteins. **(A)** Pipeline for protein intensity measurements of the seven Hox TFs (from anterior to posterior: Lab, Dfd, Scr, Antp, Ubx, AbdA and AbdB) expressed along the AP axis of the ventral nerve cord. *Upper panel:* The motoneuronal marker (FasII) in magenta was used as reference to measure Hox TF expression patterns in different embryos in a standardized manner (see Materials and Methods). *Lower panel:* Procedure of translating different fluorescence intensity measurements into standardized graphs by Fiji. **(B)** Normalized and smoothened protein measurements of Hox TFs (color code) are combined in one graph (see Materials and Methods). X-axis: relative AP position in % (normalized to length of ventral nerve cord), Y-axis = relative fluorescence intensity measurement (normalized to max. intensity). **(C)** Normalized and smoothened single cell mRNA measurements of *Hox* gene expression (color code) arranged along the inferred AP position (See Figure 1C, Materials and Methods). **(D)** Scatter plots relating principal component scores on PC3 and PC4 to AP position. PCA was performed using only cell from the *twit^low^* cluster, as in Figure S2. PCA was performed including (left panel) or excluding (right panel) *Hox* genes to rule out any qualitative biases created by targeted *HoxSeq*. p-values were computed for the hypothesis that the true correlation is different from 0 using a fisher transform of correlation coefficients. **(F)** Heatmap depicting the expression of homeodomain encoding genes (rows) across n= 758 single *twit^low^* motoneurons (columns). Normalized expression levels are color-coded in green. Rows and columns are arranged by *Hox* gene expression from anterior (A) to posterior (P) (see Figure 2), inferred position is color coded in magenta. Multi-color code indicates individual homeo-clusters as defined in Figure 1C. **(G)** The homeodomain TF Vvl and Mirr identified in ZiNB-WAVE analysis was investigated for the localisation along the DV axis of the ventral nerve cord (highlighted in white) of an early stage 17 *Drosophila* embryo. *Lower panel:* Zoom on the expression of Vvl and Mirr in the Ubx^+^ (magenta) abdominal segment 2 (A2).

**Supplementary Figure 5:**
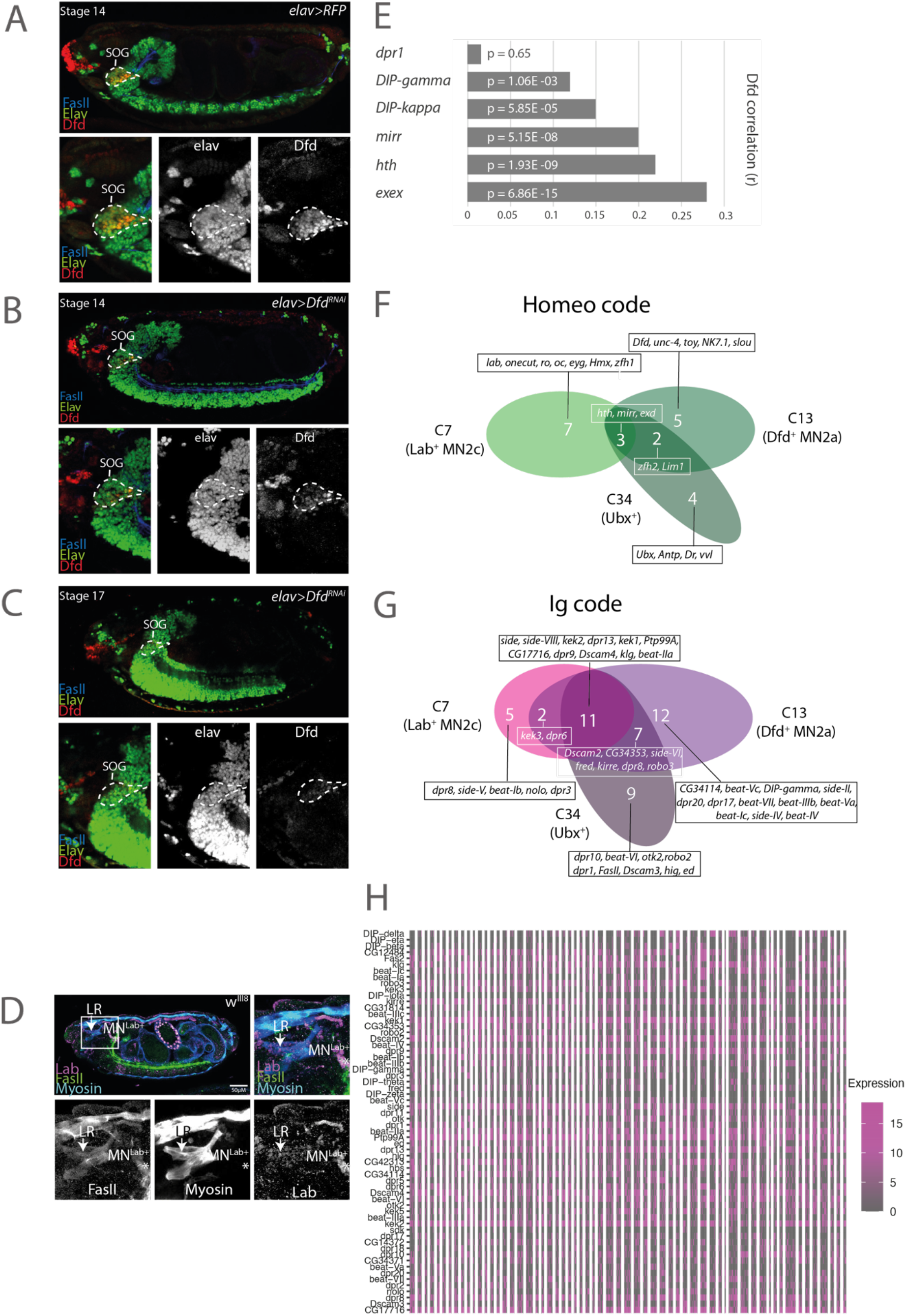
Characterisation of the Dfd expressing MN2a motoneuron. **(A)** Representative confocal image of a stage 14 *Drosophila* embryo, which depicts the lateral view of the Dfd expression pattern in the subesophageal ganglion (SOG, white circle) in *elav>RFP* control animals. **(B)** Representative confocal picture of a stage 14 embryos under *Dfd* knockdown conditions (*elav>Dfd^RNAi^*), showing a strong reduction of Dfd neuronal expression at that stage. **(C)** Representative confocal image of a stage 17 embryo displaying full depletion of *Dfd* under knockdown conditions in neurons. **(D)** The correlation between selected Ig and homeodomain encoding genes and *Dfd* expression was investigated by Pearson correlation. The homeodomain TFs encoding genes *exex, mirr* and *hth* and the Ig encoding genes *DIP-kappa* and *DIP-gamma* are significantly correlated with *Dfd* expression, while *dpr1* is not correlated with *Dfd*. p-values were computed for the hypothesis that the true correlation is different from 0 using a fisher transform of correlation coefficients. **(E-F)** Venn diagram displays the homeo-code (E) or Ig-code (F) in three different Homeodomain clusters (see cluster analysis Figure 1 C) each of the homeo-clusters is associated with a different Hox gene. **(G)** Heatmap depicting the expression of Immunoglobulin genes (rows) across n= 758 single *twit^low^* motoneurons (columns). Rows and columns are arranged by hierarchical clustering. Normalized expression levels are color-coded in magenta.

